# Macrophages foster adaptive anti-tumor immunity by ZEB1-dependent cytotoxic T cell chemoattraction

**DOI:** 10.1101/2024.02.26.582102

**Authors:** Kathrin Fuchs, Isabell Armstark, Ruthger van Roey, Yussuf Hajjaj, Elisabetta D’Avanzo, Renato Liguori, Fulvia Ferrazzi, Jochen Ackermann, Lukas Amon, Marwin Alfredo, Maria Faas, Julian Hübner, Markus H Hoffmann, Gerhard Krönke, Christoph Becker, Diana Dudziak, Falk Nimmerjahn, Simone Brabletz, Marc P. Stemmler, Thomas Brabletz, Harald Schuhwerk

## Abstract

Tumor-associated macrophages (TAMs) shape the tumor microenvironment (TME) and exert a decisive impact on anti-tumor immunity. Understanding TAM function is therefore critical to understand anti-tumor immune responses and to design immunotherapies. Here, we describe the transcription factor ZEB1, a well-known driver of epithelial-to-mesenchymal transition, as an intrinsic regulator of TAM function in adaptive anti-tumor immunity. By combining cell type-specific deletion of Zeb1 with syngeneic models of colorectal and pancreatic cancer, we discovered an unexpected function of ZEB1 in the TAM-mediated control of T cell trafficking. ZEB1 supports secretion of a subset of chemokines including CCL2 and CCL22 by promoting their transcription and translation as well as by safeguarding protein processing. ZEB1 thereby elevates cytotoxic T cell (CTL) recruitment *in vitro* and *in vivo* and fosters immunosurveillance during tumor as well as lung metastatic outgrowth. Our study spotlights ZEB1 as a crucial facilitator of adaptive anti-tumor immunity and uncovers a potential therapeutic window of opportunity for cytokine-guided enhancement of CTL infiltration into tumors and metastases.

## Introduction

Innate immunity comprises the first line of defense against cancer cells and initiates the adaptive immune response of tumor antigen-specific T cells, ideally eliciting the rejection of tumors (Hargadon 2020, Raskov et al. 2021, Falcomata et al. 2023). Hence, cytotoxic T cells (CTLs) are fundamental executors of anti-tumor and -metastasis defense (Tallon de Lara et al. 2022). However, in cancer, this defense is notoriously inefficient due to immunosuppression and/or -evasion, consistently rendering CTL infiltration a prognostic factor for survival (Maimela et al. 2019, Miksch et al. 2019, van der Leun et al. 2020, Pages et al. 2018).

Immunomodulation is the consequence of the co-evolution between tumor cells and non-malignant cells (George and Levine 2021), eventually constituting the ‘tumor microenvironment’ (TME) in solid cancers, a complex composition of extracellular matrix, fibroblasts, endothelial and immune cells (Raskov et al. 2021, Quante et al. 2013). Tumor-associated macrophages (TAMs) are one of the most abundant innate immune cell types in the TME (Pinto et al. 2019, Guo et al. 2022, Falcomata et al. 2023). Macrophages/TAMs polarize between “inflammatory” and “alternative” states – the two extreme poles of a broad spectrum of phenotypes which can either have anti- or pro-tumorigenic effects (Mantovani et al. 2022, Storz 2023). One major role of TAMs is the recruitment of immune cells such as monocytes, neutrophils and T cells via secreted chemokines (Takeya and Komohara 2016). Direct anti-tumor effects include induction of tumor cell apoptosis and phagocytosis (Chang et al. 2001, Cao et al. 2022). On the other hand, pro-tumor TAMs can promote survival, proliferation and epithelial-to-mesenchymal transition (EMT) of cancer cells by secretion of cytokines and growth factors (Haque et al. 2019, Sarode et al. 2020). Thus, macrophages/TAMs are plastic immune cells with a broad repertoire of functions that require delicate regulation in the TME.

The partial activation of the developmental process EMT in cancer cells is well-known to induce invasiveness and cellular plasticity, enabling adaptation to environmental challenges, such as posed by cancer therapies or faced during the metastatic cascade (Dongre and Weinberg 2019, Yang et al. 2020, Brabletz et al. 2021, Schuhwerk and Brabletz 2023). The core EMT-inducing transcription factor (EMT-TF) ZEB1 regulates the expression of a wide range of target genes influencing cell adhesion, differentiation, metabolism, therapy resistance, proliferation and the DNA damage response to promote plasticity and malignancy (Stemmler et al. 2019, Dongre and Weinberg 2019, Brabletz et al. 2021, Schuhwerk and Brabletz 2023).

ZEB1 is also expressed in cells of the TME (Chaffer et al. 2013, Galvan et al. 2015, Fu et al. 2019, Krebs et al. 2017, Cortes et al. 2017, Fu et al. 2020, Smita et al. 2018, Schuhwerk et al. 2022). Specifically, in cancer-associated fibroblasts (CAFs), ZEB1 was described to be crucial for fibroblast plasticity and immune infiltration supporting tumor progression (Fu et al. 2019, Bhome et al. 2022, Schuhwerk et al. 2023). Moreover, heterozygous knockout of ZEB1 was reported to impair the function of peritoneal macrophages in the context of ovarian cancer, and homozygous loss of ZEB1 in myeloid cells impaired the recovery from inflammation and the response to viral infection in mice (Cortes et al. 2017, Baasch et al. 2021, Cortes et al. 2023). However, the role of ZEB1 in TAM polarization and its impact on the TME remain elusive.

Here we employed conditional knockout mouse models and uncovered an unexpected function of ZEB1 in TAMs in suppressing metastatic lung colonization of gastrointestinal cancers. Mechanistically, we show that ZEB1 does not act as a master executor of acute polarization, but is pertinent for the secretion of TAM-derived cytokines to augment immunosurveillance.

## Results

### ZEB1 is heterogeneously expressed in macrophages in CRC and PDAC

ZEB1 is heterogeneously expressed in tumor cells and in cells of the TME (Krebs et al. 2017, Schuhwerk et al. 2022, Schuhwerk et al. 2023, Fu et al. 2019, Fu et al. 2020). Similarly, we observed heterogenous expression of ZEB1 in macrophages in publicly available single cell transcriptomes of colorectal cancer (CRC) and pancreatic cancers (Fig. 1A-D). Supporting the transcriptional data, we detected ZEB1 positive (+) CD68+ macrophages by immunofluorescence (IF) stainings of human CRC (Fig. 1E) and mouse lung metastases (Fig. 1F) formed upon tail vein injection (tvi) of pancreatic ductal adenocarcinoma (PDAC) cells (Krebs et al., 2017). In order to clarify whether ZEB1 is upregulated in macrophages, we scored the number of ZEB1+;CD68+ cells in these samples. Interestingly, tumorous tissue contained more CD68+ cells and an increased share of ZEB1+ cells among those as compared to respective healthy tissue (Fig. 1G, H). Altogether, these data demonstrate that a subset of ZEB1+ macrophages enrich in PDAC and CRC primary tumors as well as in experimental metastases.

**Figure 1:**
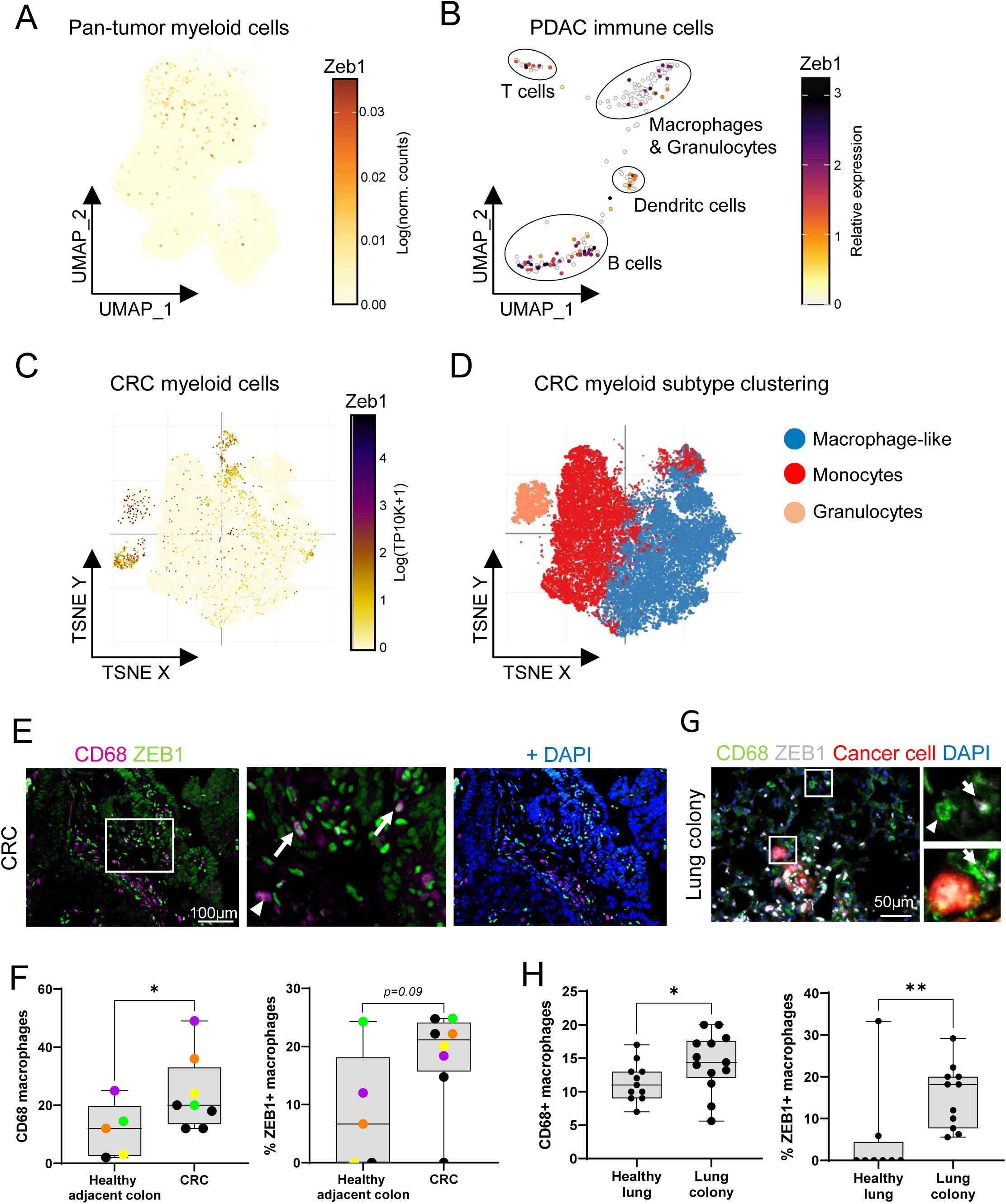
ZEB1 is heterogeneously expressed in macrophages in CRC and PDAC. **A-C**. UMAP plot visualization of Zeb1 in single cell RNA sequencing data in myeloid cells derived from human pan tumors (Cheng et al. 2021) (A), from subclustered immune cells from murine PDAC (Gabitova-Cornell et al. 2020) (B), and of myeloid cells derived from human CRC (C). **D**. Myeloid subtype clustering of cells from C, as indicated. In (C) and (D), plots and cell type annotations were retrieved from the single cell portal from the Broad Institute using the dataset from (Pelka et al. 2021). **E-F**. Representative images (E) and quantification (F) of immunofluorescence staining (IF) for CD68 and ZEB1 of human colorectal cancer (CRC) with DAPI-stained nuclei. Insets show a higher magnification. Arrows indicate CD68+;ZEB1+ cells and arrowheads CD68+;ZEB1-cells (n (healthy adjacent colon/CRC)=5/8). Color-code in (F) indicates tissue areas matched per case. **G-H**. Representative images of IF (G) and quantification (H) for CD68 and ZEB1 of murine lungs and metastatic lung colonies with DAPI-stained nuclei. Insets show a higher magnification. Arrows indicate CD68+;ZEB1+ cells and arrowheads CD68+;ZEB1-cells (n≥11; Mann-Whitney test (F, H)). *:p<0.05; **:p<0.01; ns: not significant.

### ZEB1 in macrophages is dispensable for organ development and homeostasis

To investigate the role of ZEB1 in macrophages and TAMs, we generated myeloid-specific *Zeb1*-deleted mice by combining the conditional *Zeb1* knockout mice with the ZEB1^flox^ allele (Brabletz et al. 2017) with LysM-Cre (knock-in of Cre into the *Lyz2* locus) (Clausen et al. 1999) by mating. The resulting Zeb1^flox/flox^;LysM-Cre^ki/+^ mice were designated as ‘LysM^ΔZeb1^’ and their Lyz2 wildtype littermates as ‘LysM^Ctrl^’ (Fig. S1A, B). LysM^ΔZeb1^ mice do not show any obvious phenotypic abnormalities or gross alterations in tissue architecture, as deduced from histologic analyses (Fig, S1C, D), in line with a recent study (Cortes et al. 2023). Consistently, flow cytometry of cell suspensions from colon, lung, spleen and blood from LysM^ΔZeb1^ mice revealed unchanged immune cell compositions, including macrophages (CD45+;CD11b+;F4/80+), as compared to their LysM^Ctrl^ littermates (Fig. 2A, S1E-H). To verify activity of LysM-Cre in LysM^ΔZeb1^ mice in macrophages and neutrophils, the mT/mG Cre-reporter allele was employed, which switches from ubiquitous tdTomato to GFP expression upon Cre-mediated recombination (Muzumdar et al. 2007). As expected, lymphoid and myeloid compartments in pancreata, lungs and livers of mT/mG+ LysM^ΔZeb1^ mice revealed preferential recombination in macrophages and neutrophils, as well as dendritic cells, albeit to a much lower extent in the latter (Fig. 2B-D, S2A-E). Importantly, IF analysis revealed that pancreata, colons and lungs of LysM^ΔZeb1^ mice were devoid of CD68+ZEB1+ macrophages as opposed to LysM^Ctrl^ mice (Fig. 2E, S2F). Collectively, these data show that ZEB1 is efficiently deleted from macrophages in LysM^ΔZeb1^ mice and that ZEB1 in macrophages is neither essential for organ, macrophage, neutrophil or lymphoid development nor for tissue and immune cell homeostasis.

**Figure 2:**
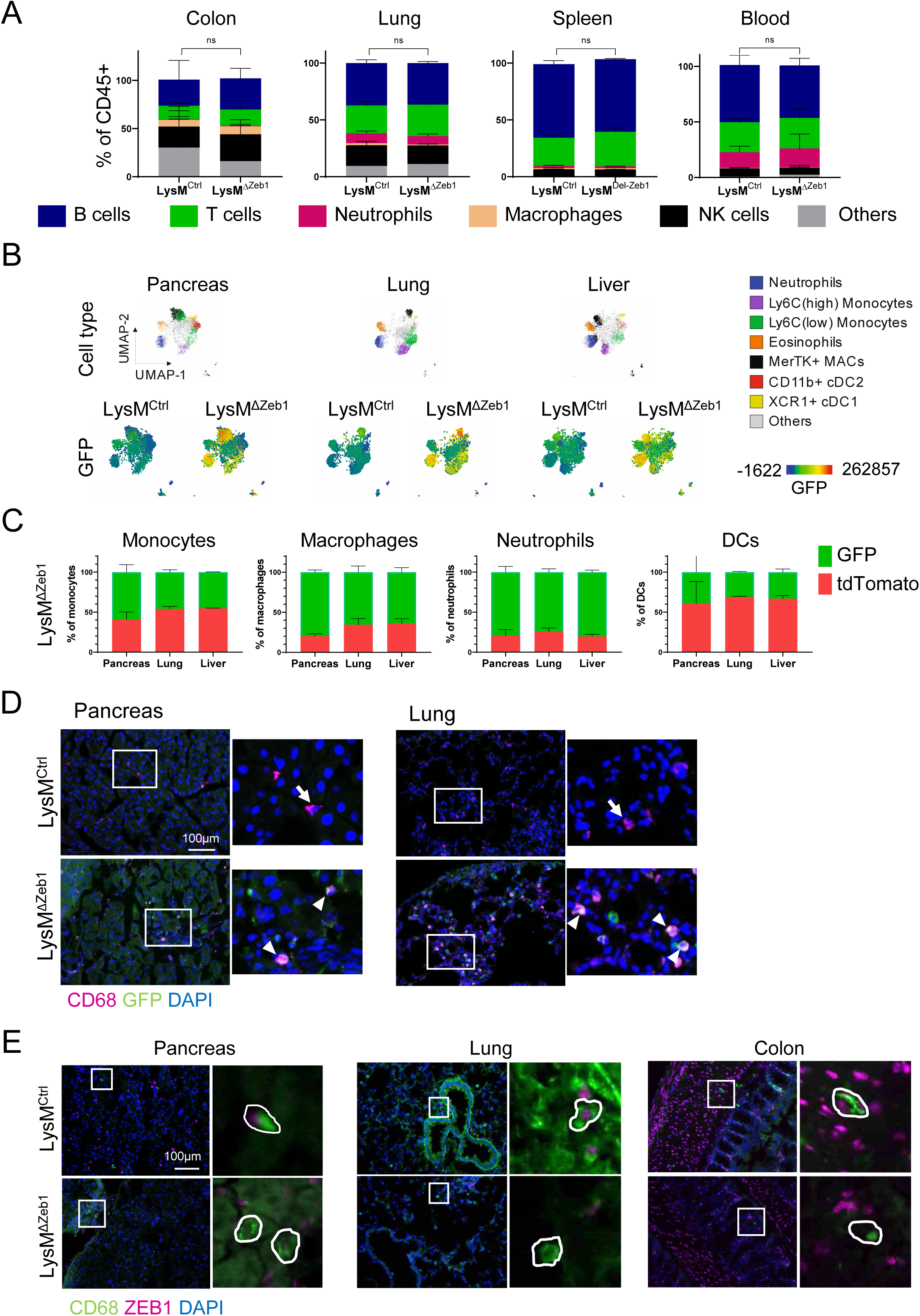
ZEB1 in LysM-expressing cells is dispensable for organogenesis and homeostasis. **A**. Percentages of B cells, T cells, neutrophils, macrophages and NK cells of CD45+ cells in organs of LysM^Ctrl^ and LysM^ΔZeb1^ mice as determined by flow cytometry (n=3; mean +SD; 2-way ANOVA). **B**. Flow cytometry UMAP clustering of cells isolated from organs of mT/mG-positive(+) LysM^Ctrl^ and LysM^ΔZeb1^ mice. GFP expression is depicted as color gradient (n=2). **C**. Percentage of tdTomato+ or GFP+ cells in immune cell subtypes of mT/mG-positive LysM^ΔZeb1^ mice (n=2; mean +SD). **D**. Representative images of CD68 IF of organs of mT/mG+ LysM^Ctrl^ and LysM^ΔZeb1^ mice with DAPI-stained nuclei. Insets show a higher magnification. Arrows show CD68+ cells. Arrowheads show CD68+;GFP+ cells. Note that the tdTomato channel is excluded. **E**. Representative images of CD68 and ZEB1 in organs of mT/mG-negative LysM^Ctrl^ and LysM^ΔZeb1^ mice with DAPI-stained nuclei (see S2F for scoring of CD68+;ZEB1+cells).

### ZEB1 in macrophages subverts tumor growth and lung colonization

To determine the impact of ZEB1 loss in macrophages on tumor growth, we first employed a syngeneic subcutaneous (s.c.) model of CRC by injecting CMT-93 cells into LysM^Ctrl^ and LysM^ΔZeb1^ mice. Interestingly, we observed failed CMT-93 tumor formation upon injection of one million cells exclusively in LysM^Ctrl^, but not in LysM^ΔZeb1^ mice (Fig.3A), suggesting that ZEB1 exerts an anti-tumor function in macrophages, which is either due to low tumorigenicity or high immunogenicity of CMT-93, or both, in LysM^Ctrl^ mice. To rule out a cell line-specific effect, we also injected highly tumorigenic MC-38 CRC cells. While no difference in tumor formation was observed upon s.c. injection of a large number of cells, MC-38 tumor formation was significantly sustained in LysM^ΔZeb1^ mice as compared to LysM^Ctrl^ mice upon reducing tumor cell numbers for engraftment (Fig. 3 B). The number of infiltrating macrophages was unchanged upon loss of ZEB1 (Fig. S3A, B). These data suggest that ZEB1 in macrophages impedes tumor outgrowth.

**Figure 3:**
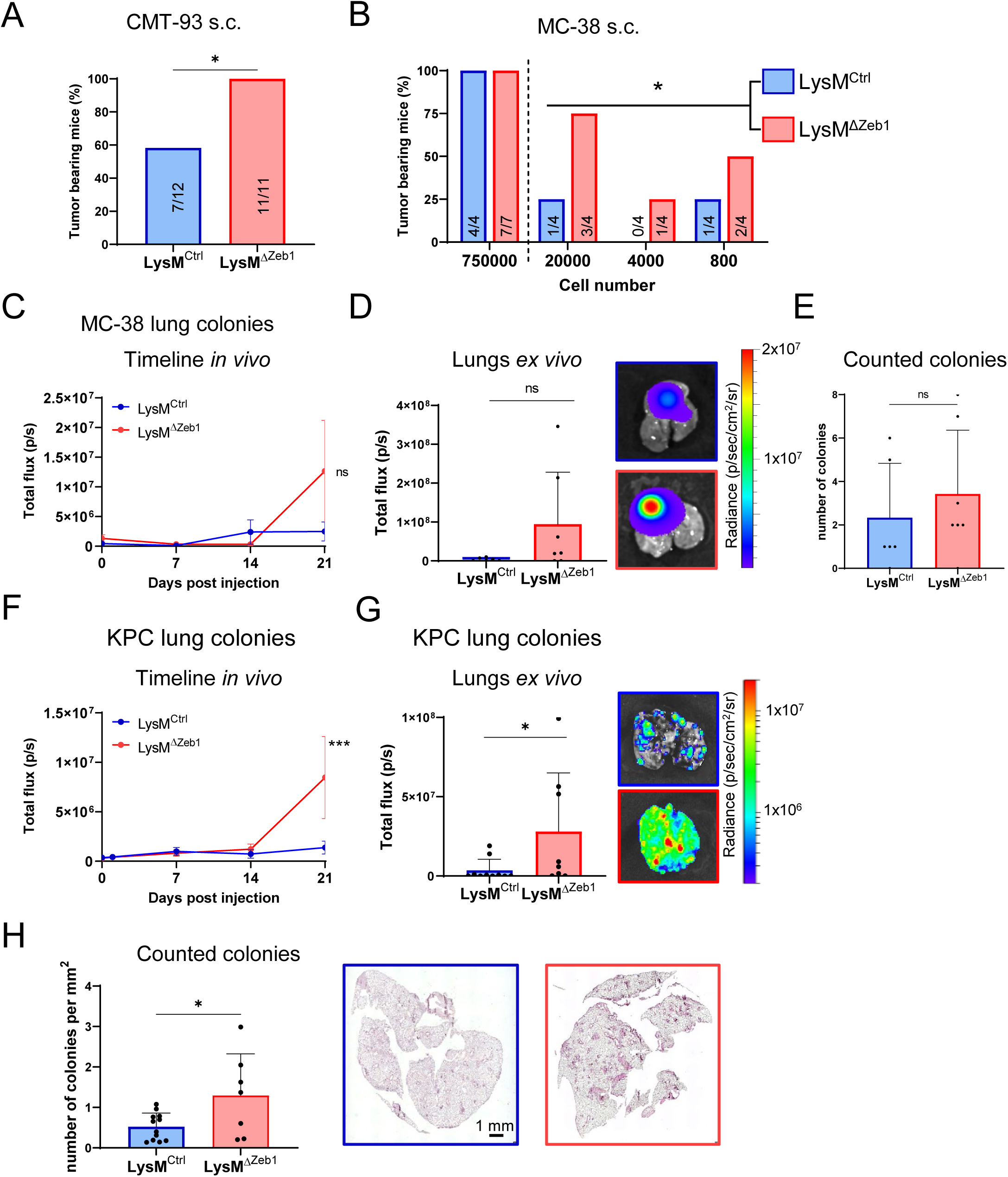
ZEB1 in macrophages subverts tumor engraftment and lung colonization. **A-B**. Percentages of tumor-bearing LysM^Ctrl^ and LysM^ΔZeb1^ mice at endpoints after s.c. injection of 1×10^6^ CMT-93 (A) and the indicated numbers of MC-38 cells (B). Number of mice (tumor-bearing/ total) is indicated in the respective bars. *: p<0.05; Fisheŕs exact test (A); (B) test for differences in stem cell frequencies using the limiting dilution assay online tool ELDA (Hu and Smyth 2009). **C**. *In vivo* BLI signal of tail vein injected MC-38 tumor cells over time in LysM^Ctrl^ (n=6) and LysM^ΔZeb1^ (n=7) mice (means ±SEM). **D**. *Ex vivo* BLI signal of lungs from mice in (C) at endpoint with representative images (means +SD) *: p<0.05. **E**. Number of counted MC-38 colonies in H&E staining of lungs of LysM^Ctrl^ (n= 6) and LysM^ΔZeb1^ (n=7) mice (means +SD). **F**. *In vivo* BLI signal of tail vein injected KPC tumor cells over time in LysM^Ctrl^ (n= 6) and LysM^ΔZeb1^ (n=7) mice (means ±SEM). **G**. *Ex vivo* BLI signal of lungs from mice in (E) at endpoint with representative images (means +SD). **H**. Number of KPC colonies and representative images of H&E-stained lungs of LysM^Ctrl^ (n=12) and LysM^ΔZeb1^ (n=7) mice (means +SD). ***: p<0.001; *:p<0.05; 2-way ANOVA (C, F); t-test (E, H); Mann-Whitney (D, G).

As CRC and PDAC frequently metastasize to the lungs, our data prompted us to clarify the potential impact of ZEB1 in macrophages on metastatic lung colonization. To achieve this, we applied an experimental metastasis model by intravenous injection of luciferase-expressing CRC (MC-38) and PDAC (KPC) cells into LysM^Ctrl^ and LysM^ΔZeb1^ mice. By longitudinal *in vivo* bioluminescence imaging (BLI) of luciferase activity, we analyzed metastatic seeding and outgrowth over time. In both models, both genotypes displayed equal BLI signals for up to 14 days post injection (dpi), suggesting similar initial metastatic seeding. Strikingly, while the signals plateaued further on in LysM^Ctrl^ mice, LysM^ΔZeb1^ mice displayed a sudden elevation of BLI signal, aggravating their metastatic burden (Fig. 3 C-H). Akin to the s.c. tumors, the number of macrophages was unchanged in metastatic lungs upon loss of ZEB1 (Fig. S3C, D). We validated the increased lung colonization of KPC cells in an additional model of mice lacking ZEB1 in monocytes and macrophages (Cx3cr1-Cre; (Yona et al. 2013); Fig. S3E). As we observed recombination in LysM-Cre mice additionally in neutrophils, we employed a model lacking ZEB1 in neutrophils (Ly6g-Cre; (Hasenberg et al. 2015) to decipher the responsible cell type, revealing a much weaker effect as compared to the LysM- and Cx3cr1-Cre strains (Fig S3F).

These findings demonstrate impaired restriction of metastatic outgrowth upon loss of ZEB1 in macrophages.

### ZEB1 is dispensable for phagocytosis but selectively promotes cytokine secretion in macrophages

To gain insights into how ZEB1 in macrophages controls tumor and metastatic outgrowth, we utilized primary bone-marrow-derived macrophages (BMDMs) from LysM^Ctrl^ and LysM^ΔZeb1^ mice. As BMDMs increase Lyz2 (LysM) expression levels during maturation within the first days *in vitro* (div) (Faust et al. 2000), we observed increasing LysM-Cre mediated recombination of the mT/mG reporter from 3 div in LysM^ΔZeb1^ BMDMs (Fig. 4A, S4A). Loss of ZEB1 in LysM^ΔZeb1^ BMDMs was verified by qPCR and western blotting of LysM-Cre-recombined GFP+ BMDMs enriched by FACS (Fig. 4A-C, S4B). FACS-enriched LysM^ΔZeb1^ BMDMs exhibited normal morphology, grew normally in cultures and did not show compensatory upregulation of other core EMT-TFs (Fig. S4B-F). We then explored an involvement of ZEB1 in archetype BMDM polarization into highly inflammatory and immunosuppressive states by stimulation with lipopolysaccharide (LPS) and IL-4, respectively. Consistent with recent literature (Cortes et al. 2023), *Zeb1* expression was upregulated in BMDMs upon LPS, but not upon IL-4 treatment (Fig. 4C, S4G), indicating that ZEB1 likely does not play a major role in IL-4-polarized, but in LPS-polarized BMDMs. Macrophages in general, but in particularly particular post LPS-dependent polarization, are efficient phagocytes and produce large amounts of inflammatory cytokines (Murray et al. 2014, Mantovani et al. 2022). As phagocytosis of fluorescent bioparticles and tumor cell debris was insignificantly - withal minimally – altered in ZEB1-deficient compared to -proficient BMDMs (Fig. 4D, E), we screened for secreted cytokines. LysM^ΔZeb1^ BMDMs secreted different amounts of various cytokines compared to LysM^Ctrl^ BMDMs in a stimulus-independent manner. Among these, particularly CCL2 and CCL22 were consistently reduced and validated in a quantitative assay (Fig. 4F, G). These data indicate an important role of ZEB1 in macrophage polarization, particularly in the secretion of the chemokines CCL2 and CCL22.

**Figure 4:**
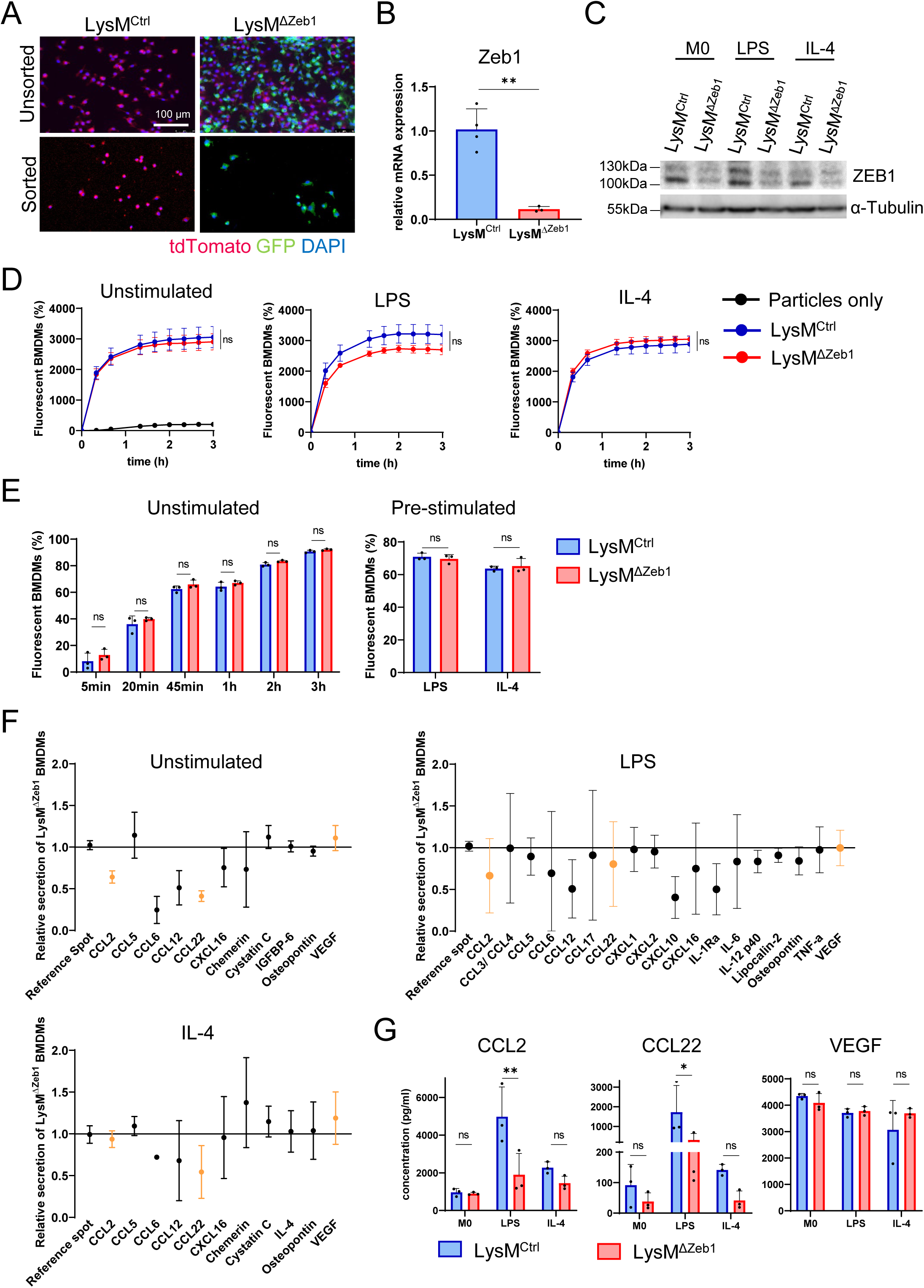
ZEB1 is dispensable for phagocytosis but selectively promotes cytokine secretion in macrophages. **A**. Representative fluorescence images of unsorted and sorted mT/mG+ LysM^Ctrl^ and LysM^ΔZeb1^ BMDMs at 8 days *in vitro* (div) with DAPI-stained nuclei. **B**. Zeb1 mRNA expression levels of sorted LysM^Ctrl^ and LysM^ΔZeb1^ BMDMs (n>3). **C**. Representative western blot of ZEB1 and α-Tubulin as loading control of unstimulated, LPS or IL-4 stimulated LysM^Ctrl^ and LysM^ΔZeb1^ BMDMs. **D**. Phagocytosis of phRodo E.coli particles by unstimulated, LPS or IL-4 pre-stimulated LysM^Ctrl^ and LysM^ΔZeb1^ BMDMs, as measured by percentage of GFP+ BMDMs (n=3 for BMDM conditions, n=1 for the particles-only condition). **E**. Time-course of phagocytosis of KPC debris by unstimulated LysM^Ctrl^ and LysM^ΔZeb1^ BMDMs or at t=20min after pre-stimulation with LPS or IL-4, as measured by percentage of Claret+ BMDMs (n=3). **F**. Secretome assays showing relative secreted molecules in supernatants of unstimulated, LPS or IL-4 pre-stimulated LysM^ΔZeb1^ BMDMs normalized to the respective LysM^Ctrl^ values (n=3). **G**. Quantification of selected cytokines using a bead-based immunoassay in LysM^Ctrl^ and LysM^ΔZeb1^ BMDM supernatants (n=3; means ±SD). *:p<0.05; **:p<0.01; ns: not significant; t-test (B, E (right)); 2-way ANOVA (E (left), G).

### ZEB1 supports inflammatory macrophage polarization and cytokine secretion by safeguarding protein biosynthesis and trafficking

We next investigated how ZEB1 supports secretion of cytokines, particularly of CCL2 and CCL22. As ZEB1 is a transcription factor, we first measured gene expression in BMDMs employing a customized qPCR array, including mostly cyto-/ and chemokines. We found moderate but insignificant differences in mRNA abundance of several cytokines, including Ccl2 and Ccl22, in LysM^Del-Zeb1^ compared with LysM^Ctrl^ BMDMs, which mostly matched the secretome data (Fig. S5A, B).

This led us to explore BMDM polarization by characterizing their general transcriptional responses to LPS and IL-4 stimulation globally by bulk RNA sequencing. As expected, LPS and IL-4 treatments changed gene expression dramatically, albeit in both genotypes. LysM^ΔZeb1^ showed more differentially expressed genes (DEGs) in response to both treatments than LysM^Ctrl^. However, 62.3% (LPS) and 55.6% (IL-4) of the DEGs were shared between the genotypes and thus apparently not dependent on ZEB1 (Fig. 5A, B). Gene Ontology (GO) term analyses identified very similar enrichments in LysM^ΔZeb1^ and LysM^Ctrl^ with respect to inflammation and the expected LPS and IL-4 responses, akin to the analysis of those DEGs shared between the genotypes (Table S1).

**Figure 5:**
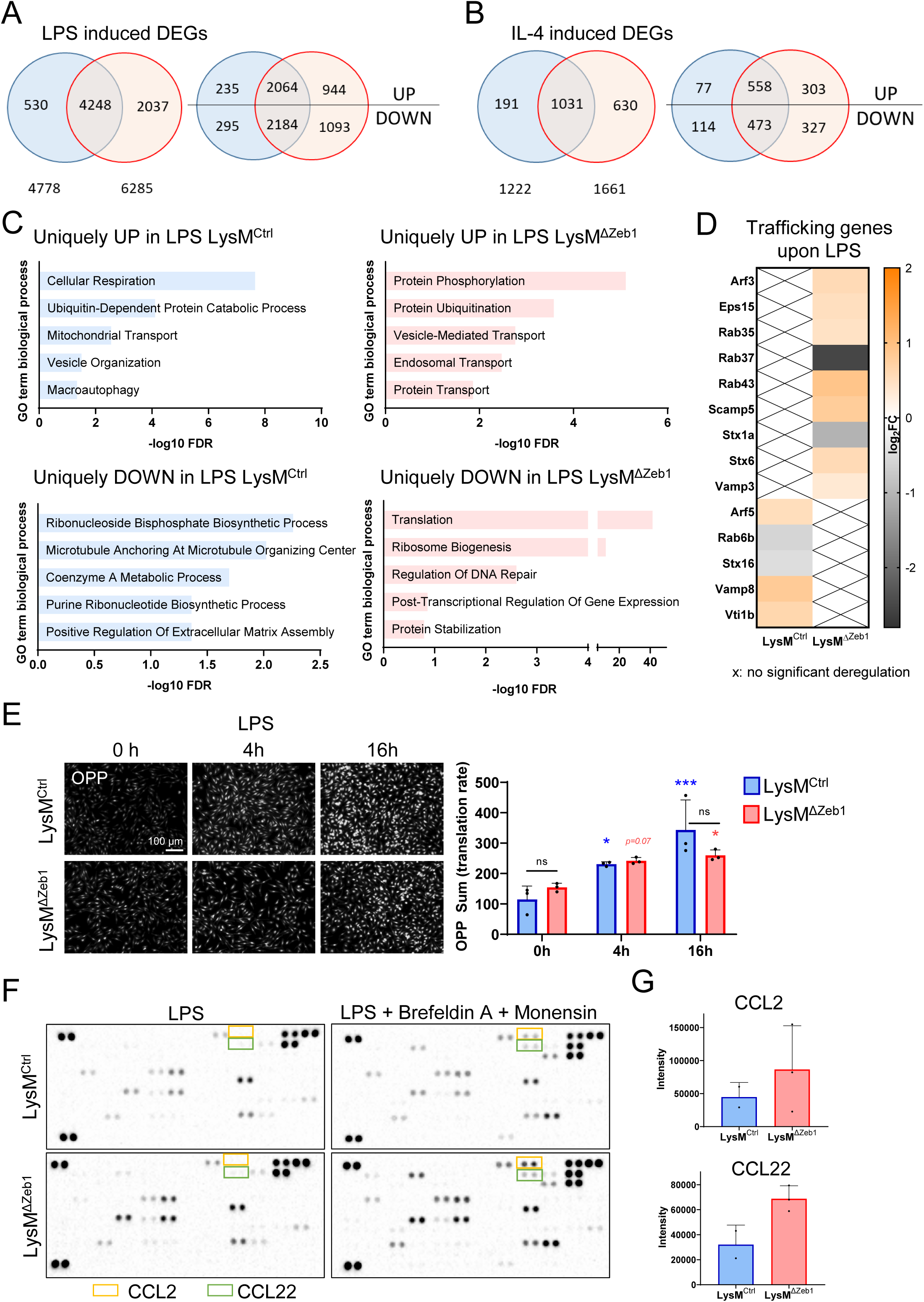
ZEB1 supports inflammatory macrophage polarization and cytokine secretion by safeguarding protein biosynthesis and trafficking. **A-B**. Venn diagrams of differentially expressed genes (DEGs) of LysM^Ctrl^ (blue) and LysM^ΔZeb1^ BMDMs (red) after stimulation with LPS (A) or IL-4 (B) compared to unstimulated in total (left) and divided in up-/downregulated DEGs (right). **C**. GO term enrichment analysis for DEGs (FDR<0.05) uniquely up- or downregulated by LysM^ΔZeb1^ BMDMs after LPS stimulation. **D**. Log_2_ fold change of expression of selected trafficking genes after LPS stimulation. X marks no significant deregulation. **E**. Representative images and quantification of OPP incorporation of LysM^Ctrl^ and LysM^ΔZeb1^ BMDMs with 0h, 4h and 16h LPS pre-stimulation (n=3; means ±SD; 2-way ANOVA). **F-G**. Representative arrays of intracellular cytokines of LysM^Ctrl^ and LysM^ΔZeb1^ BMDMs with LPS stimulation or additional Brefeldin A and Monensin treatment (F) and quantification of intracellular CCL2 and CCL22 after LPS, Brefeldin A and Monensin treatment (G; n≥2).

Intrigued by the considerable number of DEGs exclusively in LysM^Ctrl^ or LysM^ΔZeb1^ BMDMs, we performed GO term analyses on these genes (Table S2). IL-4-induced exclusive DEGs did not retrieve significant GO terms in LysM^Ctrl^ BMDMs but indicated moderately altered cell cycling in LysM^ΔZeb1^ BMDMs, consistent with the multifaceted role of ZEB1 in cell cycle control, which we addressed systematically recently (Schuhwerk et al. 2022, Schuhwerk and Brabletz 2023). In the LPS condition, LysM^Ctrl^ BMDMs showed altered cellular respiration and ribonucleotide biosynthesis, while, interestingly, protein translation, ubiquitination and transport were strongly deregulated in LysM^ΔZeb1^ BMDMs on the mRNA level, with major players in protein trafficking, such as Rab family members as well as Vamp3 being affected (Fig. 5C, D). These data indicate general polarization competence of LysM^ΔZeb1^ BMDMs concomitant to ample alterations in LysM^ΔZeb1^ BMDMs on the post-transcriptional level upon LPS treatment.

To validate the transcriptional deregulation of translation, we analyzed incorporation of O-Propargyl-Puromycin (OPP), a ‘clickable’ Puromycin analog, into nascent proteins in LysM^Ctrl^ and LysM^ΔZeb1^ BMDMs. As expected, protein synthesis increased upon LPS treatment. The full translational capacity was, however, slightly reduced in ZEB1-deficient BMDMs (Fig. 5E). In line with this, intracellular cytokines were detected to similar levels in LPS treated LysM^ΔZeb1^ compared to LysM^Ctrl^ BMDMs (Fig. 5F). Moreover, the decrease in secreted CCL2 and, particularly, CCL22 was greater than the reduction in the respective mRNA (Fig. S5C), altogether indicating a contribution from insufficient protein secretion. Indeed, blocking protein trafficking by Brefeldin A and Monensin revealed increased intracellular accumulation of cytokines in LysM^ΔZeb1^ compared to LysM^Ctrl^ BMDMs, e.g. of CCL2 and CCL22 (Fig. 5F, G). Collectively, these data indicate reduced transcription, translation and secretion of certain cytokines in ZEB1-deficient macrophages upon inflammatory polarization.

### ZEB1 in macrophages has no major direct effect on tumor cells but is essential for CCL2- and CCL22-driven recruitment of CD8+ T cells

Inflammatory cytokines can launch oncogenic signaling in tumor cells (Brabletz et al. 2021). Consistently, co-culture of BMDMs with tumor cells increased tumor cell proliferation and invasion (Fig. 6A, B), yet to the same extent in ZEB1-proficient and -deficient BMDMs. Transfer of conditioned medium from BMDMs to tumor cells had no pro-proliferative influence on them (Fig. 6C). These data suggest that macrophages do not directly influence tumor cells in a ZEB1-dependent manner.

**Figure 6:**
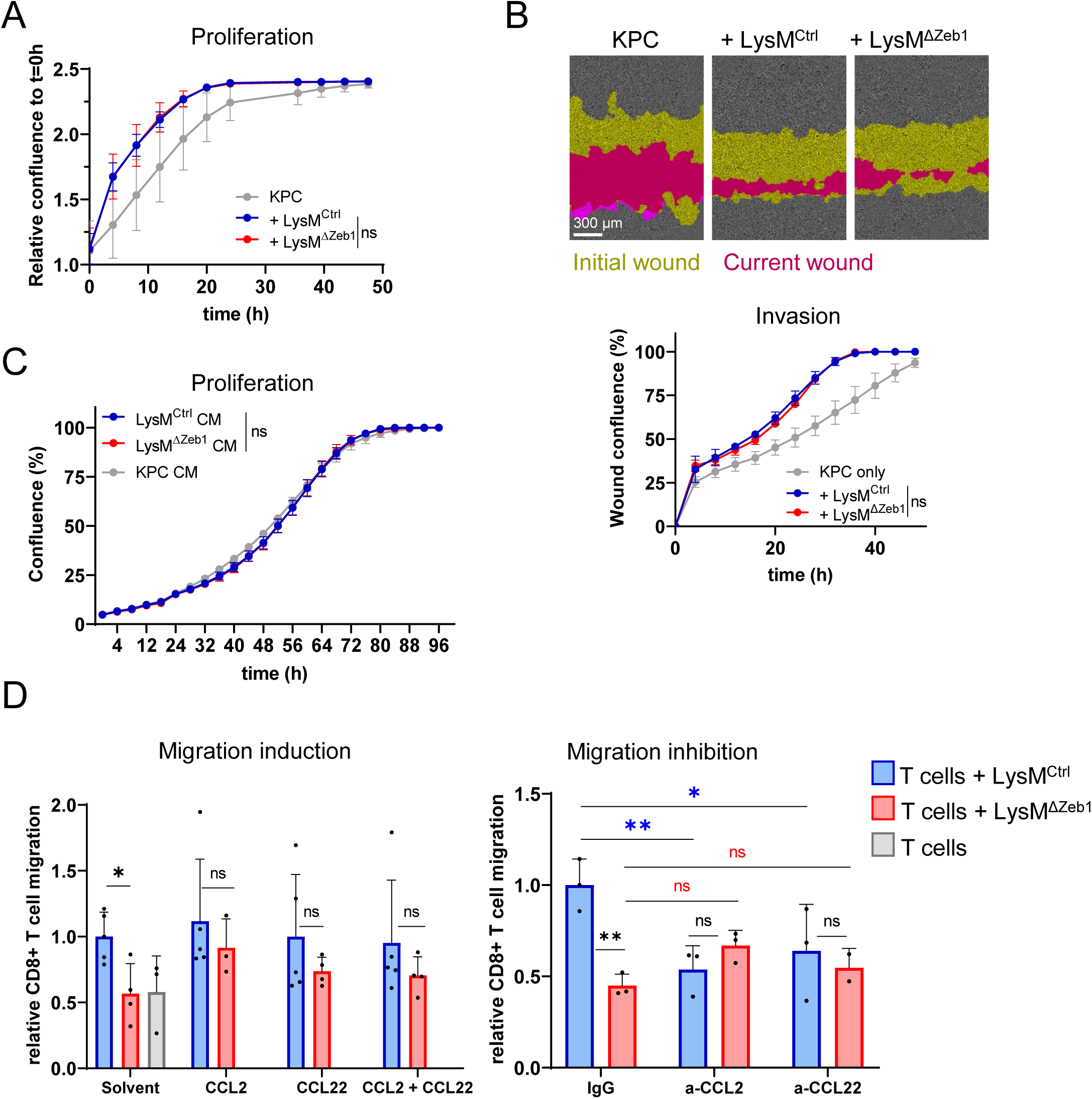
ZEB1 in macrophages has no major direct effect on tumor cells but is essential for CCL2- and CCL22-driven recruitment of CD8+ T cells. **A**. Confluence of KPC cells alone or co-cultured with LysM^Ctrl^ or LysM^ΔZeb1^ BMDMs (n=3). **B**. Representative images at t=28h and quantification over time of KPC cell invasion into a scratch wound without or with co-culture of LysM^Ctrl^ or LysM^ΔZeb1^ BMDMs (n=3). **C**. Confluence of KPC cells alone or with LysM^Ctrl^ or LysM^ΔZeb1^ BMDM conditioned medium (CM) (n=2 KPC CM, n=3 LysM^Ctrl^ and LysM^ΔZeb1^ CM). **D**. Transwell migration assay of CD8+ T cells alone or towards LysM^Ctrl^ or LysM^ΔZeb1^ BMDMs in absence or presence of recombinant CCL2 and CCL22 (left panel, n>3) or absence or presence of anti-CCL2 and anti-CCL22 antibodies (right panel, n=3). Means ±SD; *:p<0.05; **:p<0.01; ns: not significant; 2-way ANOVA.

The pleiotropic cytokines CCL2 and CCL22 are potent monocyte attractants but also recruit T cells (Okada et al. 2006, Molon et al. 2011). As monocyte influx was similar in s.c. tumors as well as metastatic lung colonies of LysM^Ctrl^ and LysM^ΔZeb1^ mice (Fig. S3 A-D), we investigated myeloid ZEB1-dependent T cell recruitment by performing T cell migration assays through narrow-pored transwells towards BMDMs. Strikingly, ZEB1-deficient BMDMs, unlike the ZEB1-proficient ones, were entirely incapable of recruiting CD8+ T cells, which was rescued by addition of recombinant CCL2 and CCL22 in amounts levelling out the concentration differences between both BMDM genotypes that were quantified before (Fig. 4G, Fig. 6D, left panel). Likewise, CCL2- and CCL22-depleting antibodies lowered CD8+ T cell recruitment in LysM^Ctrl^ BMDM co-cultures to LysM^ΔZeb1^ BMDM co-culture levels (Fig. 6D, right panel). Notably, we did not observe major differences in proliferation of CD8+ T cells upon co-culture with LysM^Ctrl^ or LysM^ΔZeb1^ BMDMs pre-stimulated with LPS or IL-4 or unstimulated (Fig. S6A). These data demonstrate that CD8+ T cells are recruited by BMDMs via CCL2 and CCL22 in a ZEB1-dependent manner.

### ZEB1 in macrophages promotes recruitment of CD8+ cells into s.c. tumors as well as during metastatic lung colonization and concomitant tumor cell killing

To clarify whether the defect in CCL2-/CCL22-dependent CD8+ T cell recruitment affects CD8+ T cell infiltration into tumors and metastatic lung colonies, we performed IHC analyses. Strikingly, we found substantially (3 times) diminished CD8+ cells in MC-38 and CMT-93 s.c. tumors, accompanied by reduced tumor cell apoptosis as marked by cleaved Caspase 3 (Fig. 7A). Underscoring a decisive influence of adaptive immunity on impeding outgrowth of CMT-93 tumor cells in LysM^Ctrl^ mice, all allografted immune-incompetent NOD-SCID gamma (NSG) mice formed tumors and were thus almost indistinguishable from LysM^ΔZeb1^ mice, whereas tumor formation failed in almost 50% of LysM^Ctrl^ mice (Fig. 7B). These data show that ZEB1 in macrophages is crucial for CD8+ cell infiltration into s.c. tumors. Similarly, in metastatic lung colonies of MC-38 cells of LysM^ΔZeb1^ mice, we observed significantly reduced tumor cell death and diminished, yet insignificant, influx of CD8+ cells (Fig. 7C). Reconciling this observation with the *in vivo* BLI dynamics of metastatic outgrowth indicating net stagnation of growth in LysM^Ctrl^ but not in LysM^ΔZeb1^ lungs, we hypothesized that CD8+ cell infiltration may precede tumor cell elimination in the lungs. To investigate this possibility and to validate the findings of the MC-38 model, we analyzed lungs by IHC at different time points during metastatic lung colonization of KPC PDAC cells. In line with our hypothesis, the number of CD8+ cells began rising at 10 dpi in colonies of LysM^Ctrl^, but not of LysM^ΔZeb1^ mice, which displayed a defect in CD8+ cell infiltration (Fig. 7D). Consequently, tumor cell death transiently peaked at 14 dpi, but to a much higher extent in LysM^Ctrl^ than in LysM^ΔZeb1^ mice. This relative difference remained until 21 dpi, despite a general reduction of tumor cell apoptosis (Fig 7E). As CCL2 and CCL22 are also chemoattractants for T helper cells (Fei et al. 2021), we scored CD4+ cells in these lung colonies. Their number increased over time, but similarly in LysM^Ctrl^ as in LysM^ΔZeb^ mice (Fig. S7).

**Figure 7:**
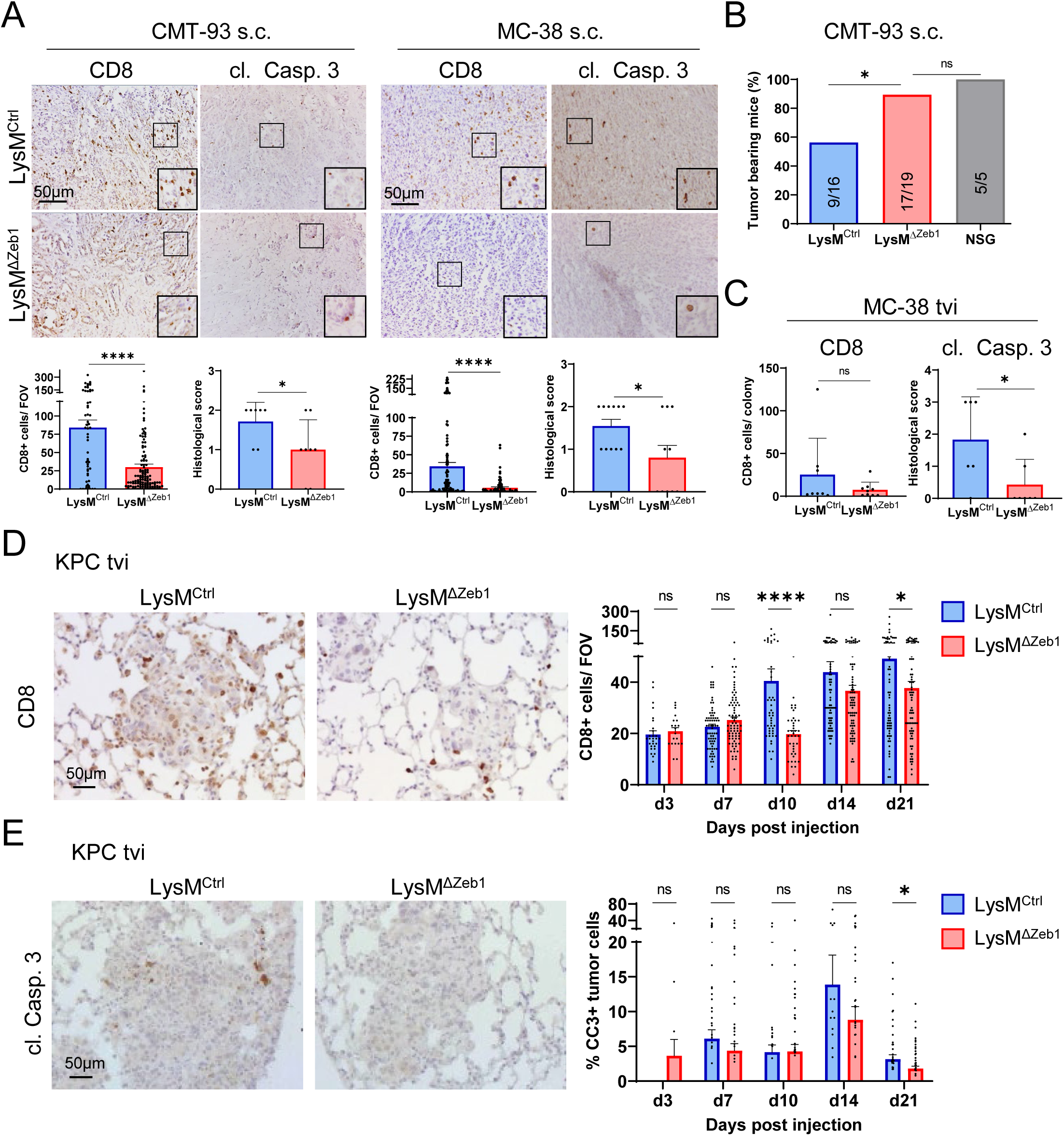
ZEB1 in macrophages promotes recruitment of CD8+ cells and concomitant tumor cell killing. **A**. Representative images and quantification of IHC for CD8 positive (+) (n>57 images of n=7 mice) and cleaved Caspase 3+ (cl. Casp. 3; n>7 mice) cells in s.c. CMT-93 and MC-38 tumors in LysM^Ctrl^ and LysM^ΔZeb1^ mice. Insets show higher magnification. **B**. Percentage of s.c. CMT-93 tumor bearing LysM^Ctrl^, LysM^ΔZeb1^ and NSG mice. Number of mice are indicated (tumor-bearing/ total). **C**. Quantification of IHC for CD8+ and cl. Casp. 3+ cells in MC-38 lung colonies in LysM^Ctrl^ and LysM^ΔZeb1^ mice (n>6). **D**. Representative images at 10 dpi and quantification over time of IHC for CD8+ cells in KPC lung colonies in LysM^Ctrl^ and LysM^ΔZeb1^ mice (n>20 images per condition). **E**. Representative images at 14dpi and quantification over time of IHC for cl. Casp. 3+ cells in KPC lung colonies in LysM^Ctrl^ and LysM^ΔZeb1^ mice (n>10 images per condition; means +SEM). ****:p<0.0001; ***:p<0.001 *:p<0.05; ns: not significant; Mann-Whitney (A); Qui-square test (B); t-test (A, C); 2-way ANOVA (D, E).

In summary, our data show that ZEB1 in macrophages promotes CD8+ cell infiltration into s.c. tumors and metastatic colonies in the lung, preceding tumor cell death to counteract tumor outgrowth.

## Discussion

We demonstrate that myeloid ZEB1 is dispensable for organogenesis and hematopoiesis, but in TAMs, it modulates the anti-tumor immune response. TAMs are mostly associated with tumor promotion, but also exhibit tumor-suppressive functions (Storz 2023, Malfitano et al. 2020). We discovered an important function of ZEB1 in macrophages to limit heterotopic tumor and lung metastatic growth by cytokine-mediated CTL recruitment in syngeneic models of two gastrointestinal tumor entities, without direct noticeable effects on tumor cells.

We show that ZEB1 does not orchestrate global macrophage polarization, but is essential for chemoattraction of CTLs. Mechanistically, ZEB1 assists inflammatory polarization, influences cytokine expression, is supporting their full translation rate and is required for their secretion. Altogether, this fine-tuning role of ZEB1 is distinct from its well-established one as a key driver of oncogenic (TGFβ-induced) EMT in tumor cells, yet adds a translationally relevant facet of ZEB1-linked cellular plasticity (Krebs et al. 2017, Caramel et al. 2018, Feldker et al. 2020, Brabletz et al. 2021, Schuhwerk and Brabletz 2023).

Consistent with our data, an ovarian cancer model revealed haploinsufficiency of ZEB1 in CCL2 secretion from peritoneal macrophages, which, unlike to what we observed, induced an EMT-like phenotype in tumor cells (Cortes et al. 2017). The discrepancy may stem from an entity-dependent effect on tumor cells and/or from heterozygous zygotic Zeb1 deletion, which may have clouded ZEB1′s specific functions in TAMs.

In agreement with our findings, the recently reported ZEB1-involving transition from an inflammatory into an immunosuppressive state of macrophages was not mainly elicited by regulation of effector genes and cytokines directly. Instead, ZEB1 controlled mitochondrial translation and autophagy, implying the involvement of intracellular membrane fusions and cargo trafficking (Cortes et al. 2023). In macrophages, defects in synthesis and processing of cytokines diminish abundance of extracellular cytokines (Murray and Stow 2014, Hughes and Nibbs 2018), as we observed in ZEB1-deficient macrophages. In our study, different Rab and Arf family members, which are crucial for the fusion of membranes during secretion (Murray and Stow 2014), were deregulated upon LPS stimulation in a ZEB1-dependent manner. This suggests that ZEB1 regulates cytokine trafficking in macrophages, expanding the described role of ZEB1 in vesicular trafficking in cancer cells (Banerjee et al. 2021, Xiao et al. 2023). Thus, cytokine secretion in macrophages appears to be supported by ZEB1 via activation of cytokine gene expression and the regulation of their posttranscriptional processing and trafficking.

ZEB1 is reportedly expressed in neutrophils, which are relevant in cancer (Scott and Omilusik 2019, Giese et al. 2019, Hsu et al. 2020), but a functional involvement has not been documented yet. As *Zeb1* deletion specifically in neutrophils had no significant tumor-promoting effect in our study, we conclude that myeloid ZEB1 exerts its anti-tumorigenic effect mostly via TAMs. In addition to macrophages, ZEB1 was reported to be important for dendritic cells, regulating their activation and production of cytokines (Smita et al. 2018), as well as for antigen cross-presentation, thereby eliciting CD8+ T cell responses (Wang et al. 2023). The role of ZEB1 therein was limiting phago-lysosomal fusions to enhance antigen export, reinforcing our notion of the importance of ZEB1 in macrophages for intracellular trafficking. Taken together, ZEB1 emerges as a multifaceted player in the adaptive immune response. We showed that ZEB1 in macrophages augments the secretion of CCL2 and CCL22 which was required for CD8+ T cell migration *in vitro*. Several studies have shown the chemotactic effect of the CCL2-CCR2 and CCL22-CCR4 axes on CTLs *in vitro* (Korbecki et al. 2020, Berencsi et al. 2011, Zhang et al. 2006, Li et al. 2020, Kondo and Takiguchi 2009) and *in vivo*, partially with an overall anti-tumor effect (Lanca et al. 2013, Okada et al. 2006, Maeda et al. 2021, Molon et al. 2011, Asai et al. 2013). In contrast, also pro-tumorigenic effects of CCL2 and CCL22 in the TME are well established, comprising e.g. monocyte attraction (Gschwandtner et al. 2019, Hao et al. 2020) as well as recruitment of T helper cells (Lim et al. 2016, Fei et al. 2021, Rapp et al. 2019, Qian et al. 2011). This was, however, unaffected in myeloid ZEB1-deficient mice and we corroborate the efficient chemotaxis of CD8+ T cells by CCL2 and CCL22 in physiologically relevant concentrations. On that account, we conclude that CD8+ T cell specific tumoricidal effects of CCL2 and CCL22 may outweigh pro-tumorigenic and immune-suppressive effects via other cell types in our models.

Notably, in clinical trials, inhibitory therapeutic targeting of the CCL2 and CCL22 axes were often unfavorable for patients (Lim et al. 2016, Fei et al. 2021, Maeda et al. 2021), reinforcing an overall anti-tumorigenic effect of CCL2 and CCL22 at least in certain tumor settings (Lehmann et al. 2017). Our data on CD8+ T cell influxes in s.c. tumors and lung metastases and other studies concordantly describing homing of CCR4+ CTLs to diseased skin or lung (Casciano et al. 2020, Spoerl et al. 2021, Mikhak et al. 2013), jointly provide an encouraging therapeutic prospect to improve T cell influx to metastasized lungs by local CCL2 or CCL22 administration.

Collectively, our study identified a new tumor-suppressive function of ZEB1 in macrophages in the TME, appeals for caution in designing anti-TAM therapies and points to a potential therapeutic window of opportunity for topical CCL2 or CCL22 supplementation in patients bearing micro-metastatic lungs.

## Methods

### Animal experiments

Animal husbandry and the experiments were approved by the committee of ethics of animal experiments of the state of Bavaria (Regierung von Unterfranken, Würzburg; TS-18/14, TS-30-2021, 55.2-DMS-2532-2-270,-2-1234) and performed according to the European Animal Welfare laws and guidelines. Animals were kept on a 12:12 hours light-dark cycle and provided with food and water ad libitum in the animal facilities of the Friedrich-Alexander University of Erlangen-Nürnberg. *Zeb1*^flox/flox^ conditional Zeb1 knockout mice, LysM-Cre, Rosa26-mT/mG, Cx3cr1-Cre and Ly6g-Cre-Tom mice have been described previously (Brabletz et al. 2017; Clausen et al. 1999; Muzumdar et al. 2007; Yona et al. 2013; Hasenberg et al. 2015). All mouse strains were kept on C57BL/6 background. *Zeb1*^flox/flox^ mice were crossed with LysM-Cre*^Ki/+^* mice to obtain LysM-Cre*^Ki/+^*;Zeb1^flox/+^ which were crossed *inter se* to obtain *Zeb1*^flox/flox^;LysM-Cre*^+/+^*(‘LysM^Ctrl^’) and LysM-Cre*^Ki/+^*;*Zeb1*^flox/flox^ (‘LysM^ΔZeb1^’) mice. *Zeb1*^flox/flox^ mice harboring either one Cx3cr1-Cre or one Ly6g-Cre-Tom allele instead of LysM-Cre were generated by breeding as described above for ‘LysM^Ctrl^’ and ‘LysM^ΔZeb1^’ mice. The Rosa26-mT/mG allele was crossed in as necessary and all mice were PCR genotyped for all alleles (see Table S3 for oligonucleotides). NOD.Cg-Prkdcscid Il2rgtm1WjI/SzJ (Nod-Scid-gamma, NSG) were bred in-house. Age-matched littermates of both sexes were used for experiments.

### Tumor cell culture and retroviral transduction

Tumor cell lines and their genetically modified derivatives (tdTomato, Luciferase, as indicated) were cultured in DMEM containing 10% FBS and 1% P/S in a humidified incubator (5% CO2) at 37 °C.

Tumor cell transduction for constitutive luciferase and tdTomato expression was carried out by ecotropic retroviral infection. The pLib_EF1A_nlstdTomato-2A-puro plasmid was generated by HiFi DNA assembly (Cell Signaling, E2621) according to the manufacturer’s instructions. In brief, PCR fragments containing pLib vector backbone (Clontech) and EF1A promoter, a cassette coding for tdTomato with an N-terminal nuclear localization sequence and lack of a stop codon and a puromycin cassette with an N-terminal self-cleaving T2A peptide were amplified by KOD polymerase (Merck, 71085) and subjected to Gibson assembly.

For retroviral transduction, 2×10^6^ PlatE cells were plated into 10 cm plates in DMEM/10% FBS, transfected with 8 µg of pLib_EF1A_LuciNeo (kindly provided by Ralf Graeser, ProQinase, Freiburg) or pLib_EF1A_nlstdTomato-2A-puro using FugeneHD (Promega). Medium was exchanged after 4-8 h. Virus-containing supernatant of PlatE cells (virus-SN) was harvested and filtered through 0.2 µm membranes after 2 days, and directly used for viral transduction of cancer cell lines. 1×10^5^ KPC and MC-38 cells were plated one day before transduction into 6-wells and transduced by replacing the medium with 2 ml of virus-SN supplemented with 8 µg/ml polybrene. After 3-4 h of incubation, the transduction medium was replaced by standard medium. Starting from the day after, cells were selected by G418 (KPC661-Luc, MC-38-Luc; 175-300 µg/ml for 7 days) or puromycin (KPC661-nTom; 1.5-1.75 µg/ml for 4 days), where appropriate. tdTomato-positive cells were enriched by FACS and expanded.

### Subcutaneous tumor allografts

Cancer cells in 100 μl PBS (containing 1×10^6^ CMT-93, 1×10^5^ KPC661 or MC-38 as indicated) were subcutaneously injected into flanks of mice. Mouse weight and tumor size were measured by calipering. Mice were sacrificed when their or their littermateś tumor reached a critical size or ulcerated.

### Lung colonization and bioluminescence imaging

100 μl PBS containing 2×10^5^ MC-38, 2×10^5^ KPC661-Luc or 2×10^5^ KPC661-nTom was injected into the tail vein of mice. Mice were monitored at least twice per week and sacrificed at indicated time points. Growth of tumors with Luciferase reporter was monitored by bioluminescence imaging. 10 min after injection of 100 μl D-Luciferin-Na-salt (25 mg/ml, PJK, 102133) in PBS s.c. in anesthetized mice, bioluminescence signal was measured in the IVIS® Spectrum In Vivo Imaging System (Perkin Elmer). Lung ex vivo bioluminescence was measured in the IVIS® system directly after sacrifice. BLI images were analyzed using the Living Image® software.

### Histology and immunohistochemistry (IHC)

Organs or tumors were collected and either fixed in 4% paraformaldehyde, and embedded in paraffin (FFPE) or cryopreserved by freshly freezing the tissue in Tissue-Tek® O.C.T.™ Compound (Sakura Finetek, 4583). Lungs were inflated with PBS or via intratracheal injection of 60% Tissue-Tek® O.C.T.™ Compound in PBS prior to FFPE or cryopreservation, respectively.

For hematoxylin and eosin (H&E) staining, FFPE slides were deparaffinized and rehydrated in Roti®-histol (4x 15 min), isopropanol (2x 5 min) and ethanol (2x 5 min 96% ethanol, 5 min 80% ethanol, 5 min 70% ethanol) and washed 5 min in deionized H_2_O. Slides were stained in 1:10 hematoxylin solution in Milipore H2O and counterstained with 0.2% eosin in 100% ethanol. Slides were dehydrated in an ethanol row (80% ethanol, 2x 100% ethanol), followed by 2x isopropanol and xylol. For mounting, Roti®-Histokitt was used.

IHC was performed as previously described (Schuhwerk et al. 2023) using the antibodies provided in table S3.

For immunofluorescence (IF), cryopreserved tissue on glass slides were permeabilized and quenched for 10 min in PBS containing 0.1M glycine and 0.1% Triton^TM^ X-100. Following 3x washing in PBS, tissues or cells were blocked in blocking buffer (3% BSA in PBS) for 30 min. Slides were incubated with primary antibodies, diluted in blocking buffer, overnight at 4 °C. The following day, slides were washed 3x in PBS and incubated with fluorophore-coupled secondary antibodies for 1 h. Details about all antibodies are provided in table S3. Following washing 3x in PBS, slides were incubated 15 min in 1μg/ml DAPI working solution. Slides were washed in 3x PBS and mounted with CitiFluor^TM^. Images were acquired using a Leica DM5500B microscope. Automated quantification of histological stainings was performed using ImageJ. Briefly, images were formatted to RGB, deconvoluted for RGB and background subtracted for thresholding of positive cells.

### Flow cytometry

After opening the abdomen and thorax, about 300 μl blood was slowly extracted from the right ventricle of the heart with an EDTA flushed syringe and added to 14 ml Hank′s solution. Samples were centrifuged for 10 min at 1400 rpm and all but 3 ml supernatant was removed. After blood harvesting, mice were perfused with 0.9% NaCl and organs collected in ice-cold PBS. Organs were minced with scissors and incubated in 10 ml digestion mix (DMEM/ F-12 containing 0.05% Collagenase D (Sigma-Aldrich, 11088858001), 0.3% Dispase II (Sigma-Aldrich, D4693) and 0.05% DNase I (Sigma-Aldrich, 10104159001)) for 25 mins at 37 °C with gentle agitation. Digestion was stopped by adding 30 ml ice-cold PBS, the samples were filtered through a 70 μm cell strainer and centrifuged at 1200 rpm at 4 °C for 5 min. The cell pellet was resuspended in 3 ml ACK Lysis buffer. After 3 min, erythrocyte lysis was stopped with 27 ml PBS and the samples centrifuged for 5 min at 1200 rpm at 4 °C. 1×10^6^ cells were blocked in 50μl TruStain FcX™ PLUS in PBS for 15 min a RT. 50 μl of 2x FACS antibody mix was added and incubated for 20 min at 4 °C in the dark. Details about all antibodies are provided in table S3. Following centrifugation for 5 min at 1200 rpm at 4 °C, cell pellets were resuspended in 500 μl FACS buffer. Samples were analyzed in the CytoFlex Analyzer (Beckman Coulter). Data was analyzed using the CytExpert or Kaluza software (Beckman Coulter).

For flow cytometry cross-tissue UMAP clustering, samples were stained in 50µL of FACS buffer (PBS+2%FCS) containing purified FC blocking antibodies (clone 2.4G2 and clone 9E9) as well as the monoclonal biotin coupled antibodies for a total of 20 minutes at 4°C. Next, samples were washed three times with FACS buffer. Subsequently, cells were stained with monoclonal fluorescently-labeled antibodies and streptavidin-BUV496. Details about all antibodies are provided in table S3. After three washing steps, samples were supplemented with FACS buffer containing DAPI in a final concentration of 50 ng/mL and analyzed using a LSRFortessa™ SORP (BD). Data was analyzed using FlowJo (BD). 5000 (myeloid panel) or 2150 (lymphoid panel) cells per tissue were randomly selected from the pool of CD45+ cells, combined to form an analysis sample and UMAP was performed. The individual samples were again separated by tissue and the populations identified by gating.

### Bone marrow-derived macrophage (BMDM) isolation and culture

On day 0, bone marrow of mouse femur and tibia bones was flushed with 10 ml PBS using a 26G cannula. Following centrifugation for 5 min at 1200 rpm, the cell pellet was resuspended in 3 ml ACK Lysis buffer. After 3 min, erythrocyte lysis was stopped with 27 ml PBS and cell suspension was filtered using a 70 μm cell strainer. Following centrifugation for 5 min at 1200 rpm, the cell pellet was resuspended in 10 ml BMDM medium (DMEM containing 10% FBS, 10% L-929 supernatant and 1% P/S), plated in a 10 cm petri dish and cultured in a humidified incubator (5% CO2) at 37 °C. On day 1, the supernatant containing BMDMs was collected and centrifuged for 5 min at 1200 rpm and BMDMs were seeded in BMDM medium (5-6×10^6^, 0.3×10^6^, 0.1×10^6^ and 0.015×10^6^cells/ well or dish in 10cm dishes, 6-well, 24-well and 96-well plates, respectively. On day 3, additional BMDM medium was added. When required, cells were sorted according to mT/mG fluorophores on day 4 by FACS. To this end, BMDMs were detached by incubation with CellStripper (Corning, 25-056-CI) for 15 min at 37 °C, followed by 5 min at 4 °C. After sorting (MoFlo^TM^ XDP, Beckman Coulter), cells were plated as described above. When required, BMDMs were stimulated with 10 ng/ml LPS (Sigma-Aldrich, L2630) or 20 ng/ ml IL-4 (BioLegend, 574304) for 24 h, unless indicated otherwise.

For BMDM confluence assays, BMDMs were cultured in the IncuCyte ZOOM® incubator (Sartorius) in a 24-well plate from day 1 on. Images of each well were acquired automatically every 4 hours by the IncuCyte ZOOM® on-board camera and then analyzed using the IncuCyte ZOOM® 2018A software.

For BMDM cell death assay, BMDMs were cultured with 1:1000 SYTOX™ Green Ready Flow™ Reagent (Invitrogen, R37168) in the IncuCyte ZOOM® incubator in a 96-well plate from day 6 on. Cells were cultured and monitored using IncuCyte. Image acquisition and analysis was performed as described above.

### Phagocytosis assays

For bioparticle assays, 100 ng/ml pHrodo Green E. coli BioParticles™ (Thermo Fisher Scientific, P35366) were added to BMDMs, on day 7. Cells were cultured in the IncuCyte ZOOM® incubator. Image acquisition and analysis was performed as described above.

For phagocytosis of necrotic cell debris, KPC661-Luc cells were labeled with CellVue® Claret (Sigma-Aldrich, MIDICLARET) following manufactureŕs instructions. 4×10^6^ cells/ ml were incubated in a water bath at 60 °C for 30 min to create necrotic bait cells. BMDMs (5 div) were stimulated with 10 ng/ml LPS or 20 ng/ ml IL-4. On day 6, stimuli were removed and 100 μl of necrotic cells were added to each well for indicated time points. BMDMs were washed 4x with ice-cold PBS and incubated 15 min at 37 °C with CellStripper. Detached cells were collected in FACS tubes, centrifuged at 1200 rpm for 5 min and resuspended in 500 μl FACS buffer. Data was acquired in the CytoFlex analyzer (Beckman Coulter) and analyzed using the CytExpert or Kaluza software (Beckman Coulter).

### Immunofluorescence staining and protein translation labeling via OPP

For mT/mG reporter visualization in cultured BMDMs, BMDMs were washed, fixed with 4% PFA for 15 min and permeabilized and quenched for 10 min in PBS containing 0.1M glycine and 0.1% Triton^TM^ X-100. Following three washes in PBS, slides were incubated 15 min in 1μg/ml DAPI in PBS. After three washes in PBS, slides were mounted on glass slides using CitiFluor^TM^.

Protein translation was detected with the “Click-iT™ Plus OPP Alexa Fluor™ 488 Proteinsynthese-Assay-Kit” (Thermo Fisher Scientific, C10456) according to manufactureŕs instructions. Briefly, BMDMs in 96-well plates were stimulated with LPS for the indicated timepoints followed by 2h of 10µM OPP incubation before fixation and quenching as described above. Click reaction was carried out for 1 hour at room temperature. Subsequently, IF staining for ZEB1 was performed by incubating primary antibody over night at 4°C and secondary antibody for 1 h at room temperature (details about antibodies are provided in table S3). DAPI was used as counterstain (1-2 µg/ml) and cells were kept in PBS until image acquisition. 20 images per well were taken using the EVOS M7000 microscope (Invitrogen). Images were analyzed using CellProfiler (Kamentsky et al. 2011), employing previously optimized pipelines (Schuhwerk et al. 2022). Remaining strongly Zeb1-positive cells in LysM^ΔZeb1^ BMDM cultures were excluded from the analysis and a minimum of 5000 cells were analyzed per condition in 3 independently isolated LysM^Ctrl^ and LysM^ΔZeb1^ BMDM lines.

### Analysis of gene expression

For RNA isolation, cDNA synthesis and quantitative reverse transcriptase PCR, cells were washed with PBS and lysed with 350 μl RLT Plus from the RNeasy Plus Mini Kit (QIAGEN, 74136) before RNA isolation following manufactureŕs instructions. cDNA was synthesized using the RevertAid First Strand cDNA synthesis Kit (Thermo Fisher Scientific, K1622) according to manufactureŕs instructions.

qRT-PCR was performed in triplicates in 384-well plates using primers and Roche universal probe library (Roche, 04869877001) with TaqMan^TM^ Universal MasterMix II (Thermo Fisher, 4440044) and LightCycler® 480 II (Roche). Alternatively, mRNA expression levels were measured with the RT^2^ Profiler PCR Array with a custom designed panel (QIAGEN, 3445261) using the Power SYBR™ Green PCR Master Mix (QIAGEN, 4367659) according to manufactureŕs instructions. Details about primers for qPCR are provided in table S3 Bulk RNA sequencing of BMDM RNA was performed by Novogene using poly(A) enrichment library preparation and paired-end sequencing (PE150).

### Bulk RNA sequencing analysis

Bulk RNA sequencing of BMDM RNA (n=3 per condition) was performed by Novogene using poly(A) enrichment library preparation protocol and paired-end sequencing (PE150). The preprocessing of raw RNA-seq data (FASTQ files) was performed using the nf-core RNA-seq pipeline v.3.8.11 (Ewels et al. 2020). In particular, reads were adapter- and quality-trimmed using Trim Galore v.0.6.7 (https://github.com/FelixKrueger/TrimGalore). The reads were then mapped to the Ensembl mouse genome assembly GRCm39 (release 107) using STAR v.2.7.10a (Dobin et al. 2013). For transcript-level read counting, Salmon v.1.5.2 (Patro et al. 2017) was employed, relying on the Ensembl gene annotation file release 107. To generate gene-level counts, Salmon’s transcript-level quantification files were processed using the R-package tximport v.1.22 (Soneson et al. 2015) within R v.4.0.3. Differential expression analysis for the RNA-seq data was performed using the DESeq2 package v.1.34.0 (Love et al. 2014). As DESeq2 design formula ∼*litter* + *group* was utilized, where the *group* factor was created by merging together genotype (LysM^Ctrl^/LysM^ΔZeb1^) and treatment (LPS/IL-4). Fold change shrinkage was performed relying on the “ashr” method (Stephens 2017), made available within the DESeq2 package. GO term analysis was performed using Enrichr (Kuleshov et al. 2016). For this, for all significantly differentially expressed genes (DEGs) from the indicated comparisons (adjusted p value: FDR<0.05) were used and the -log10 (FDR) values plotted with GraphPad Prism. The numbers of identified DEGs for the indicated comparisons that were used for GO term analyses are indicated in the Venn-like diagram in Fig. 5A, B.

### Analysis of single-cell RNA sequencing data

For pan-cancer single cell sequencing analysis of myeloid cells, the online tool of (Cheng et al. 2021) was utilized (http://panmyeloid.cancer-pku.cn). The dataset ‘Pan_cancer scanorama_corrected’ was selected and filtered for ‘Macro’, ‘Mono’, ‘Monolike’ and ‘Myleoid’ clusters. The Zeb1 feature plot (on UMAP clusters) was directly retrieved from the portal.

For murine pancreatic cancers, the FASTQ files of (Gabitova-Cornell et al. 2020) were aligned to the reference transcriptome CellRanger 7.1.0 (Zheng et al. 2017) and subjected to further processing using Seurat 5.0., involving quality control (number of genes detected in each cell > 300; total number of molecules detected per cell > 500; mitochondrial read count ratio cutoff <25%; Haemoglobin ratio <10%), normalization of counts using Seurat′s ‘SCTransform’, integration of the three datasets using Seurat′s ‘RPCAIntegration’, and clustering. Next, DECORDER annotation was used (Chijimatsu et al. 2022) to annotate the individual cells, of which immune cells were UMAP sub-clustered. ‘FindAllMarkers’ (top 50 DEGs of a given cluster versus all other cell clusters) allowed annotation of individual immune cell types/clusters (B cells: CD2+;Bkl+;Fcmr+, T cells: Gata2+;Itk+; dendritic cells: Dcstamp+;Slamf9+, macrophages/granulocytes: Tlr4+;Ccrl2+). The feature plot in Fig. 1B was created using ‘SCPubr’ (https://enblacar.github.io/SCpubr-book/closing_remarks/Citation.html).

For CRC single cell sequencing analysis of myeloid cells, the built-in online tool from the Broad Institutés ‘single cell portal’ (https://singlecell.broadinstitute.org/single_cell) was used to explore the dataset from (Pelka et al. 2021). Plots and cell type annotations were directly retrieved from the portal. For myeloid subtype clustering, cell types were filtered for ‘myeloid’ cells (in Fig. 1C, Zeb1 feature plot), and, among those, for ‘macrophage-like’, ‘monocytes’ and ‘granulocytes’ (in Fig. 1D for comparison), using the respective available filters.

### Protein isolation

Protein was isolated as previously described (Schuhwerk et al. 2022). Briefly, cells were washed twice with PBS. Triple lysis buffer containing 150 mM NaCl, 50 mM Tris-HCl pH 8.0, 0.5% Na-Desoxycholate (w/v), 0.1% SDS (v/v), 1% NP40 (v/v), 1 mM PMSF, 1x complete protease inhibitor cocktail (Roche, 04693132001), 1x PhosStop (Roche, 4906837001) was added, cells scraped off and transferred to a 1.5 ml tube for cell lysis before clearance by centrifugation and protein concentration measurement using the Pierce™ BCA Protein Assay Kit (Thermo Fisher Scientific, 23225) following manufactureŕs instructions and the FLUOstar Omega reader. After SDS-PAGE, proteins were transferred onto nitrocellulose membrane (Roth, 4685.1) by wet blot transfer. Membranes were subjected to immunoblotting using the antibodies defined in Table Material and ECL-based signal revelation according to the manufactureŕs instructions (Western Lightning Plus-ECL, NEL103001EA, Perkin Elmer).

### Cytokine protein arrays

The Proteome Profiler Mouse XL Cytokine Array (R&D systems; ARY028) was used according to manufactureŕs instructions in order to detect secreted proteins in 500µl cell culture supernatant or intracellular proteins in 100 µg protein of cell lysates. Where indicated, cells were pre-treated with Protein-Transport-Inhibitor-Cocktail (eBioscience™, 00-4980-03) for 6 h. Quantitative analysis of secreted proteins was performed using a LEGENDplex^TM^ Custom Mouse 14-plex Panel (BioLegend).

### Co-culture assays and conditioned medium transfer

For proliferation assays, BMDMs (7 div) and KPC661-Luc cells were seeded in a 1:3 ratio to a total number of 8×10^4^ cells/ well into a 24 well plate. For the invasion assay, BMDMs (7 div) and KPC661-Luc cells were seeded in a 1:3 ratio to a total number of 1.6×10^4^ cells/ 96-well. After 4h, monolayers were scratched with the IncuCyte® 96-Well Scratch Woundmaker. Cells were washed twice with pre-warmed culture medium and 50 μl 8 mg/ ml matrigel was added to each well. Following incubation in a humidified incubator (5% CO2) at 37 °C for 30 min, BMDM medium was replenished. Cells were cultured and monitored using IncuCyte. Image acquisition and analysis was performed as described above.

For assays involving conditioned medium transfer, 200 Luciferase-expressing KPC661 cells in 200 μl/ well were plated in a 96 well plate. Conditioned medium from BMDMs or KPC661-Luc cells was added every other day by removal of 100 μl medium and addition of 100 μl conditioned medium. Cells were cultured and monitored using IncuCyte. Image acquisition and analysis was performed as described above.

### T cell isolation, enrichment of CD8+ T cells and T cell-involving assays

Mouse spleens were smashed through a 70 μm cell strainer. Following centrifugation at 1200 rpm for 5 min at 4 °C, the spleenocyte pellet was resuspended in 3 ml ACK lysis buffer for 3 min. To stop erythrocyte lysis, 27 ml ice cold PBS was added and the samples centrifuged at 1200 rpm for 5 min at 4 °C. Spleenocytes were resuspended in 3 ml MojoSort™ buffer and enriched for CD8+ T cells using the MojoSort™ Mouse CD8 T Cell Isolation Kit (BioLegend, 480035), following manufactureŕs instructions.

For the T cell attraction assays, 500 μl of double concentrated cytokines or antibodies in BMDM medium were added to BMDMs (5 div). Final concentrations of cytokines were 1.93 ng/ ml CCL2 (R&D Systems, 479-JE-050) and 0.18 ng/ ml CCL22 (R&D Systems, 439-MD-025) and of antibodies 5 μg/ml IgG (Diagenode, C15410206), anti-CCL2 (Novus Biologicals, NBP1-07035SS) or anti-CCL22 (abcam, ab124768), respectively. 5×10^5^ isolated T cells in BMDM medium with 0.1% FBS were seeded in the upper chamber of transwells with 3 μm pore size (Greiner, 662630) placed in each well and incubated for 24 h at 37 °C in a humidified incubator (5% CO2). The semi-adherent T cells were harvested by harsh pipetting and centrifuged at 1200 rpm for 10 min. T cells were resuspended in 50 μl ice-cold PBS and counted.

For the T cell activation assay, isolated T cells were labeled with CellVue® Claret Labeling following manufactureŕs instructions. 3 μg/ml anti-CD3 (Thermo Fisher Scientific, 16-0032-82) and 5 μg/ ml anti-CD28 (Thermo Fisher Scientific, 16-0281-82) antibodies were added to 2×10^6^ T cells/ ml in RPMI medium. 2×10^5^ T cells/ well were added to BMDMs (6 div) and incubated for 3 days at 37 °C in a humidified incubator (5% CO2). After centrifugation at 1500 rpm at 4°C for 10 min, T cells were resuspended in 200 μl ice-cold FACS buffer and transferred to a fresh 96-well plate. T cells were washed with ice-cold FACS buffer. Fluorescence data was acquired in the CytoFlex analyzer (Beckman Coulter) and analyzed using the CytExpert or Kaluza software (Beckman Coulter).

### Data availability

RNA-seq data will be deposited at GEO to be publicly available as of the date of publication after peer-review. Likewise, all original code will be deposited at Zenodo and will be publicly available as of the date of publication after peer-review. Additional information required to reanalyze the data reported in this paper is available upon request.

### Statistical analysis

Statistical analysis was performed using GraphPad Prism software (version 9.3.1) and within the provided R packages or cited online tools. Data is depicted as indicated. The n-numbers represent the number of biological replicates in each group. Statistical significance was assessed as indicated, depending on assay-specific sampling as well as type and distribution of the obtained data.

## Contributions

Conceptualization, KF, TB, HS; Methodology and Validation, KF, MPS, MHH, GK, HS; Software KF, RvR, YH, RL, FF, LA, HS; Formal Analysis, KF, RvR, YH, HS; Investigation, KF, IA, MA, MF, JA, JH, LA, ED, HS; Resources, SB, RL, FF, YH, RvR, FN, DD, CB, MHH, MPS, TB, HS; Writing – Original Draft, KF, HS; Writing – Review & Editing, KF, SB, MPS, TB, HS; Supervision, TB, HS; Funding Acquisition, TB, HS.

## Acknowledgements

We are grateful for Eva Bauer, Britta Schlund and Friederike Gräbner for excellent technical assistance. We thank Uwe Appelt and Markus Mroz for excellent FACS support. We also thank Elisabeth Naschberger for infrastructural support. We are also grateful for all colleagues in the Franz-Penzoldt-Center and the Preclinical Imaging Platform Erlangen for excellent animal work support.

---

**Figure S1:**
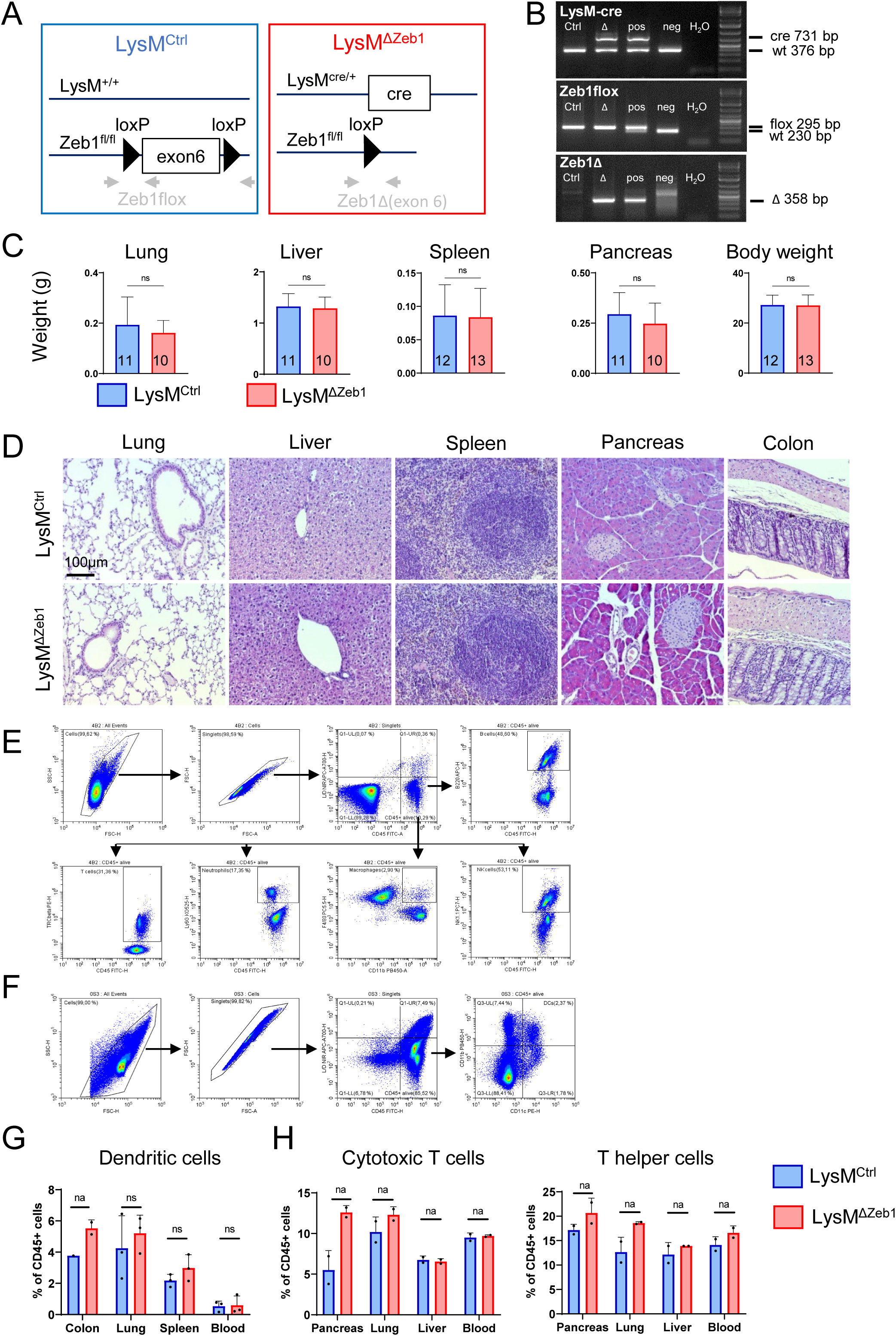
Targeting of *Zeb1* in LysM-expressing cells does not cause major phenotypic abnormalities. **A**. Schematic of the *Zeb1* targeting strategy in LysM-positive cells. Grey arrows mark primer annealing sites for genotyping. **B**. PCR genotyping of DNA isolated from LysM^Ctrl^ (Ctrl) and LysM^ΔZeb1^ (Δ) mice. wt = wild type, Δ= deleted, pos = positive control, neg = negative Ctrl, bp = base pairs. **C**. Weight of organs and animals of LysM^Ctrl^ and LysM^ΔZeb1^ mice (n indicated in figure; means +SD; t-test). **D**. Representative images of H&E stained LysM^Ctrl^ and LysM^ΔZeb1^ organs. **E**. Gating strategy for flow cytometry of immune cell subtypes in organs and blood of LysM^Ctrl^ and LysM^ΔZeb1^ mice (referring to Fig. 2A). **F**. Gating strategy for flow cytometry of dendritic cells in organs and blood of LysM^Ctrl^ and LysM^ΔZeb1^ mice (referring to Fig. S1G). **G-H.** Percentage of immune cells of LysM^Ctrl^ and LysM^ΔZeb1^ organs, as determined by flow cytometry (n(LysM^Ctrl^ / LysM^ΔZeb1^)= 1/ 2 colon; n=3 other organs (G); n=2 (H); mean +SD; t-test). ns: not significant.

**Figure S2:**
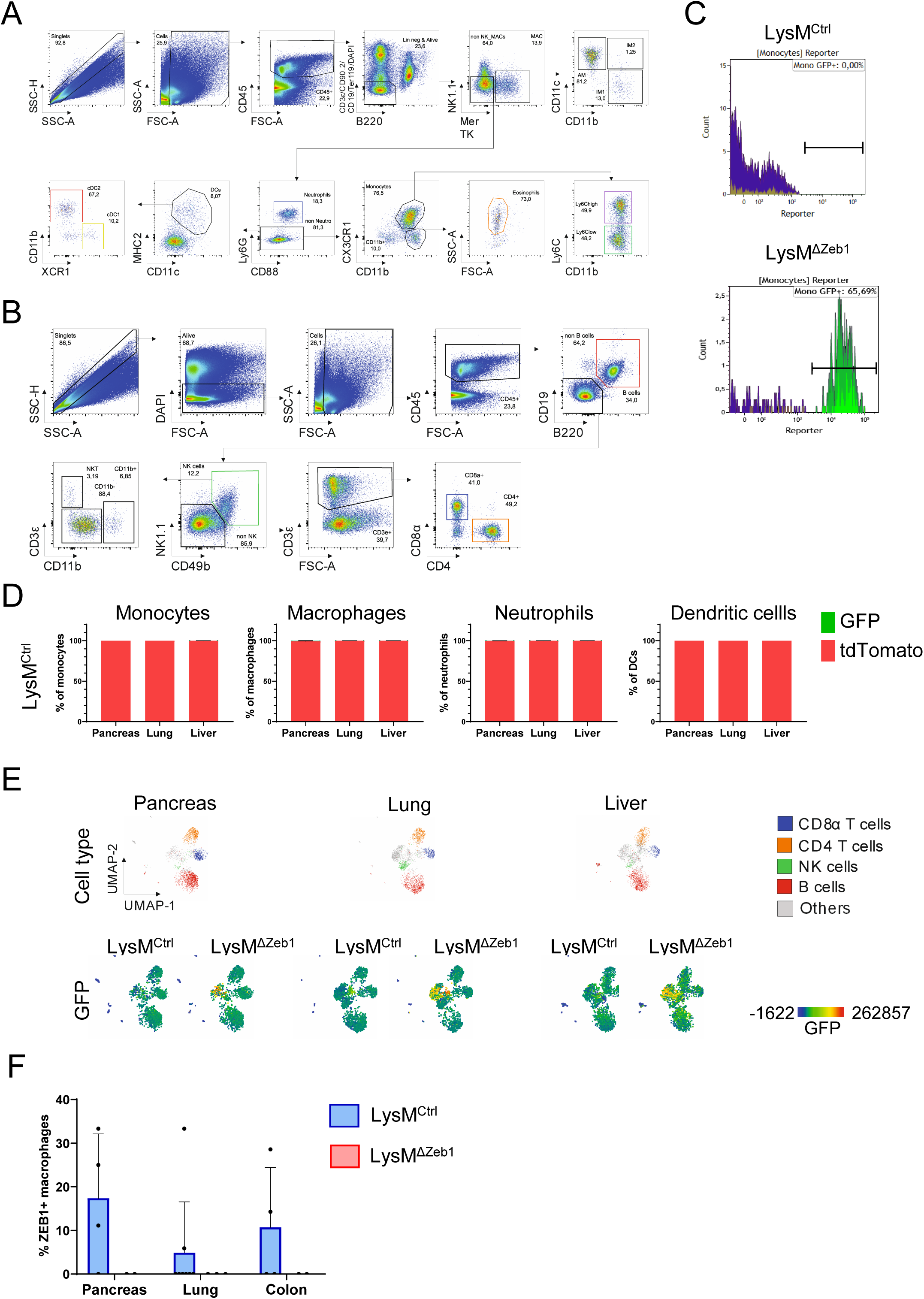
Validation of cell type specific LysM-Cre activity and ZEB1 loss in macrophages. **A-B**. Gating strategy for flow cytometry of myeloid (A; referring to Fig. 2B, C, S2D) and lymphoid (B; referring to Fig. S1H and S2E) cells isolated from mT/mG+ LysM^Ctrl^ and LysM^ΔZeb1^ mice. **C**. Representative histogram of GFP intensities in flow cytometry of cells isolated from mT/mG+ LysM^Ctrl^ and LysM^ΔZeb1^ mice used for Fig. 2C and S2D after gating according to Fig. S2A or B. **D**. Percentage of tdTomato+ or GFP+ cells in immune cell subtypes of mT/mG-positive LysM^ΔZeb1^ (n=2; mean +SD). **E**. Flow cytometry UMAP clustered cells isolated from organs of mT/mG+ LysM^Ctrl^ and LysM^ΔZeb1^ mice. GFP expression is depicted as color gradient (n=2 each). **F**. Scoring of CD68+;ZEB1+cells from IF stained tissue from mT/mG-negative LysM^Ctrl^ and LysM^ΔZeb1^ mice from Fig. 2E.

**Figure S3:**
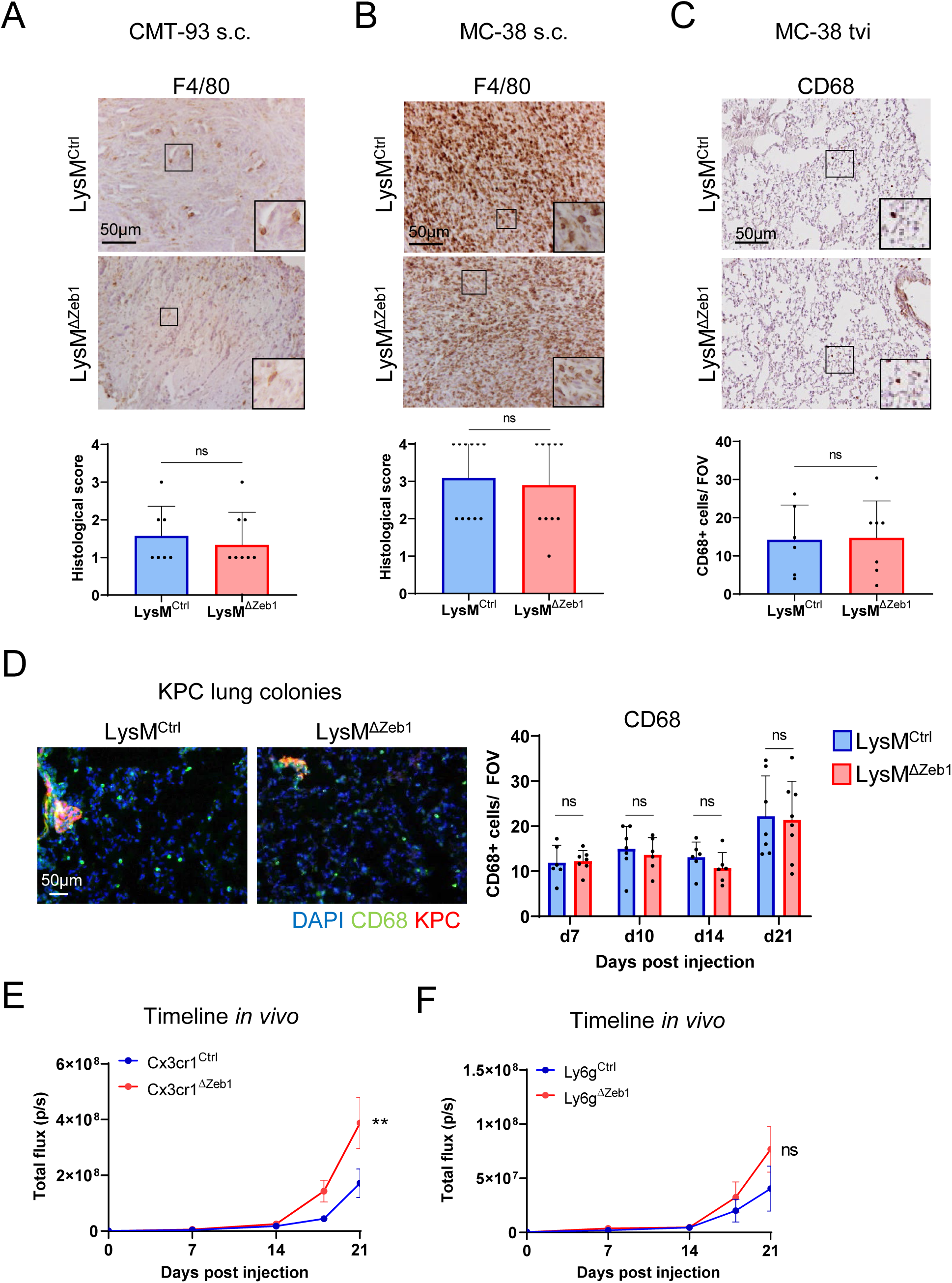
Loss of ZEB1 does not affect intratumoral macrophage infiltration in LysM-Cre mice and moderately enhances lung colonization in Cx3cr1-Cre and Ly6g-Cre mice. **A-D**. Representative images and quantifications of IHC or IF stainings for F4/80 and CD68 of s.c. CMT-93 (A), s.c. MC-38 (B) tumors and MC-38 (C) and KPC (D) lung colonies in LysM^Ctrl^ and LysM^ΔZeb1^ mice. Insets show higher magnification. **E-F**. *In vivo* BLI signal of tail vein injected KPC tumor cells over time in Cx3cr1^Ctrl^ (n=6) and Cx3cr1^ΔZeb1^ (n=12) mice (E) or Ly-6g^Ctrl^ (n= 9) and Ly-6g^ΔZeb1^ (n=7) mice (F; means ±SEM; **:p<0.01; 2-way ANOVA).

**Figure S4:**
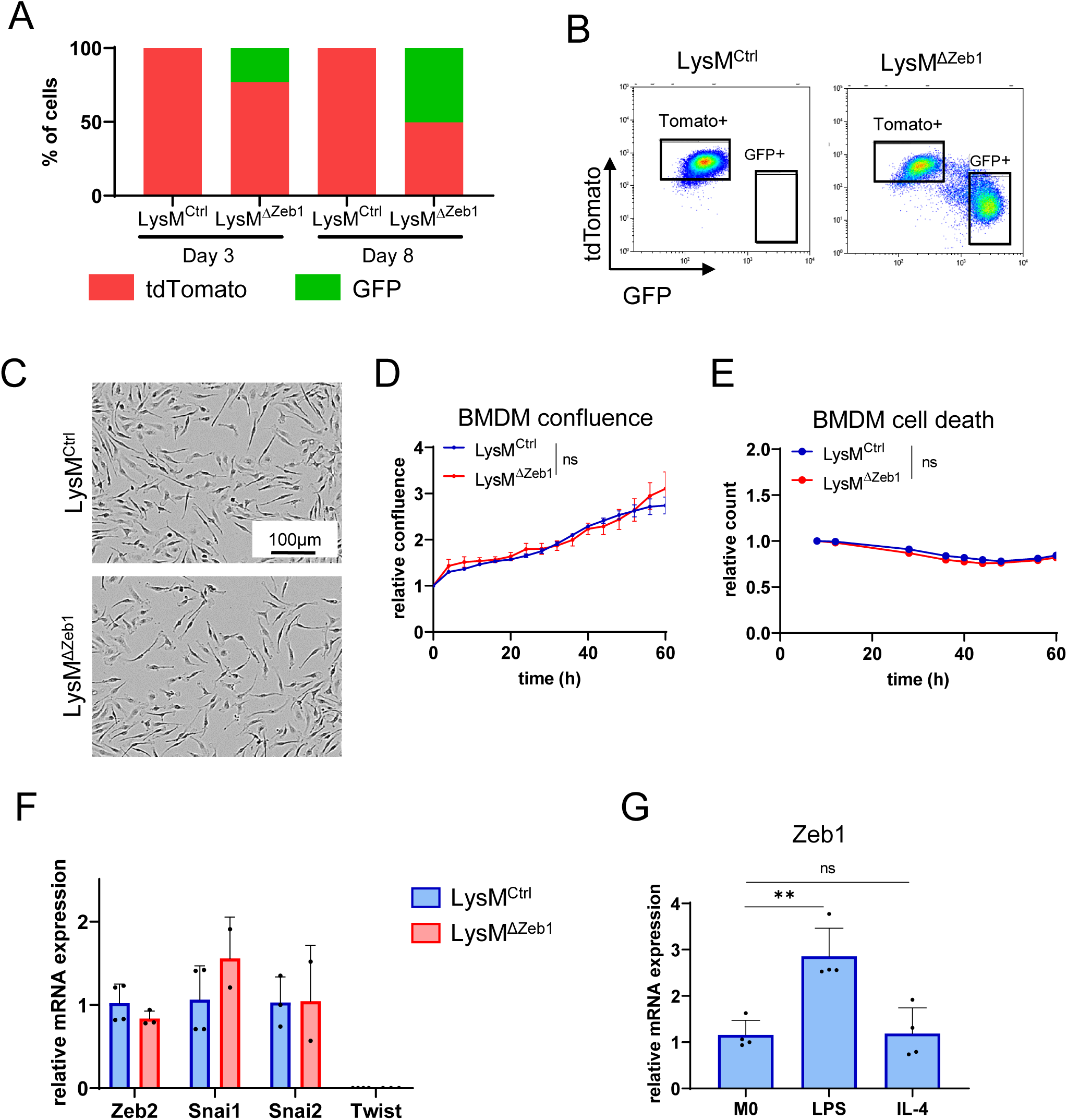
ZEB1 is dispensable for BMDM cultivation and upregulated upon LPS stimulation. **A**. Flow cytometry of LysM^Ctrl^ and LysM^ΔZeb1^ BMDMs with mT/mG reporter at 3 and 8 div (n>1.15*10^5^ cells). **B**. Representative gating for sorting LysM^Ctrl^ and LysM^ΔZeb1^ BMDMs using the mT/mG reporter. **C**. Representative bright field images of cultured LysM^Ctrl^ and LysM^ΔZeb1^ BMDMs. **D**. Relative confluence of LysM^Ctrl^ and LysM^ΔZeb1^ BMDMs over time (n=3). **E**. Relative cell death of LysM^Ctrl^ and LysM^ΔZeb1^ BMDMs as determined by SYTOX^TM^ uptake (n=3). **F**. Relative expression levels of indicated mRNAs of sorted LysM^Ctrl^ and LysM^ΔZeb1^ BMDMs (n≥2). **G**. Relative Zeb1 mRNA expression levels of sorted LysM^Ctrl^ and LysM^ΔZeb1^ BMDMs after LPS or IL-4 stimulation (n=4). Mean ±SD; **:p<0.01; 2-way ANOVA (D, E); t-test (F), 1-way ANOVA (G).

**Figure S5:**
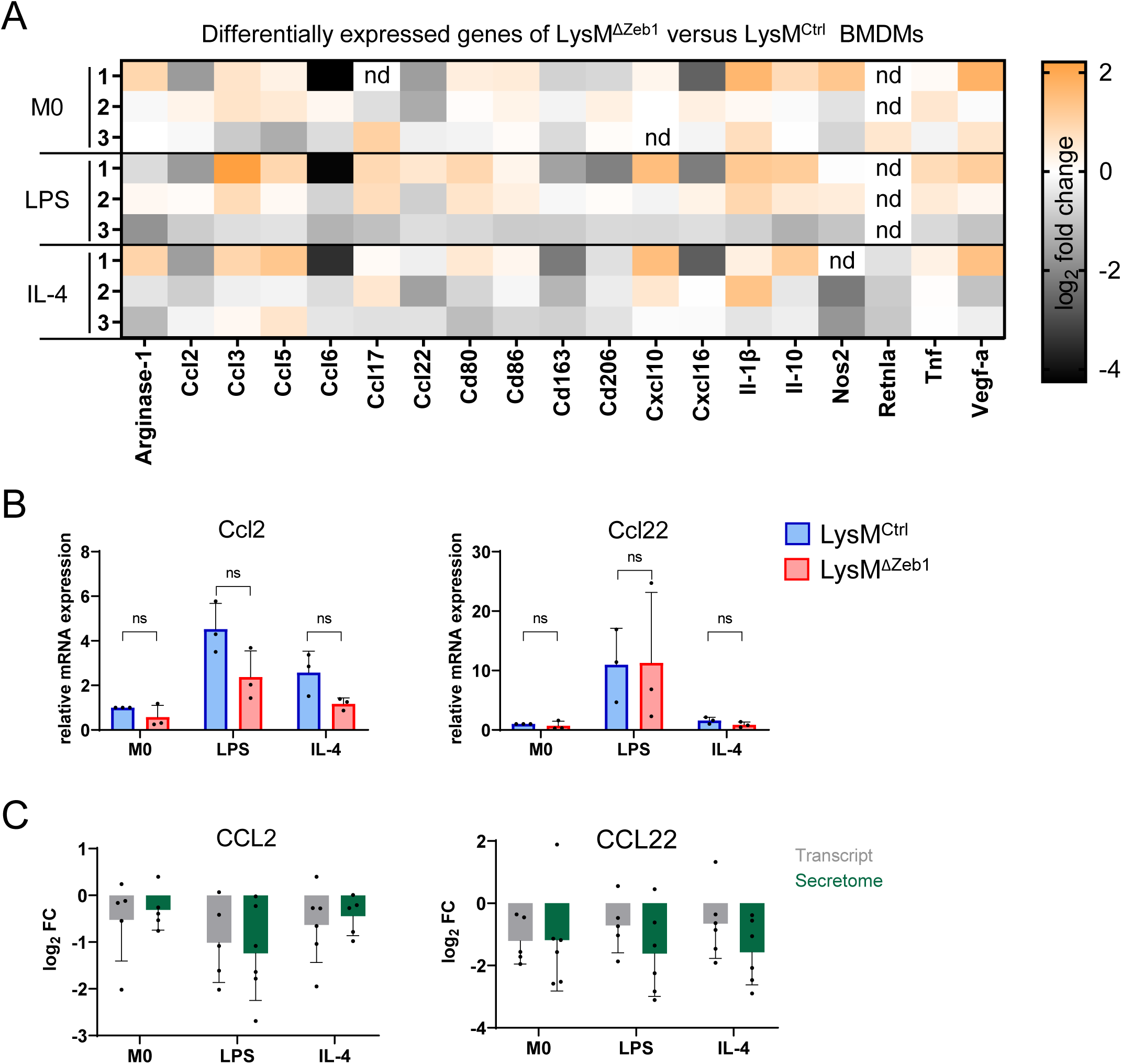
Expression and secretion of selected polarization markers cytokines in LysM^Ctrl^ and LysM^Del-Zeb1^ BMDMs. **A**. Differentially expressed genes in unstimulated or LPS or IL-4 stimulated LysM^ΔZeb1^ compared to LysM^Ctrl^ BMDMs as measured by a customized RT2 array and depicted in log_2_ fold change of expression. nd marks non-detectable mRNA levels. All transcripts were normalized to Gapdh (n=3). **B**. Relative mRNA expression of Ccl2 and Ccl22 in LysM^Ctrl^ and LysM^ΔZeb1^ BMDMs (n=3; means ±SD; 2-way ANOVA). **C**. Comparison of transcript and secretome alterations of Ccl2 and Ccl22 in LysM^ΔZeb1^ compared to LysM^Ctrl^ BMDMs (n≥5; means ±SD).

**Figure S6:**
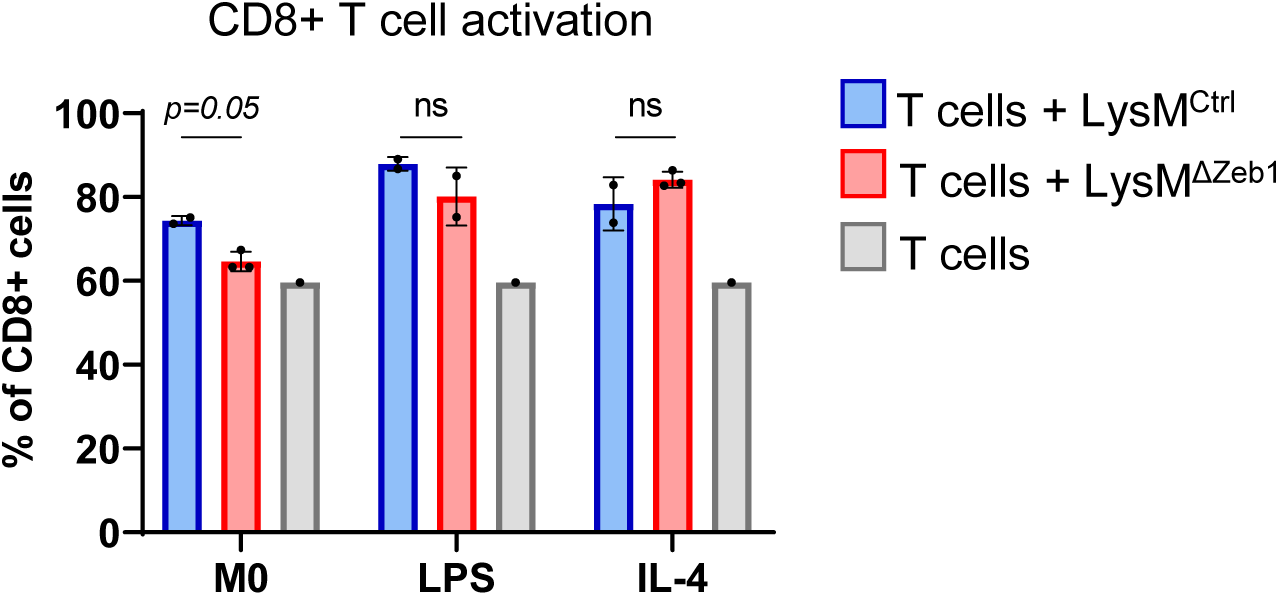
ZEB1 in macrophages has no major effect on CD8+ T cell proliferation. Activated CD8+ T cells as percentage of total CD8+ T cells in absence (n=1) or presence of M0, LPS or IL-4 pre-stimulated LysM^Ctrl^ (n=2) or LysM^ΔZeb1^ (n=3) BMDMs (means +SD; 2-way ANOVA).

**Figure S7:**
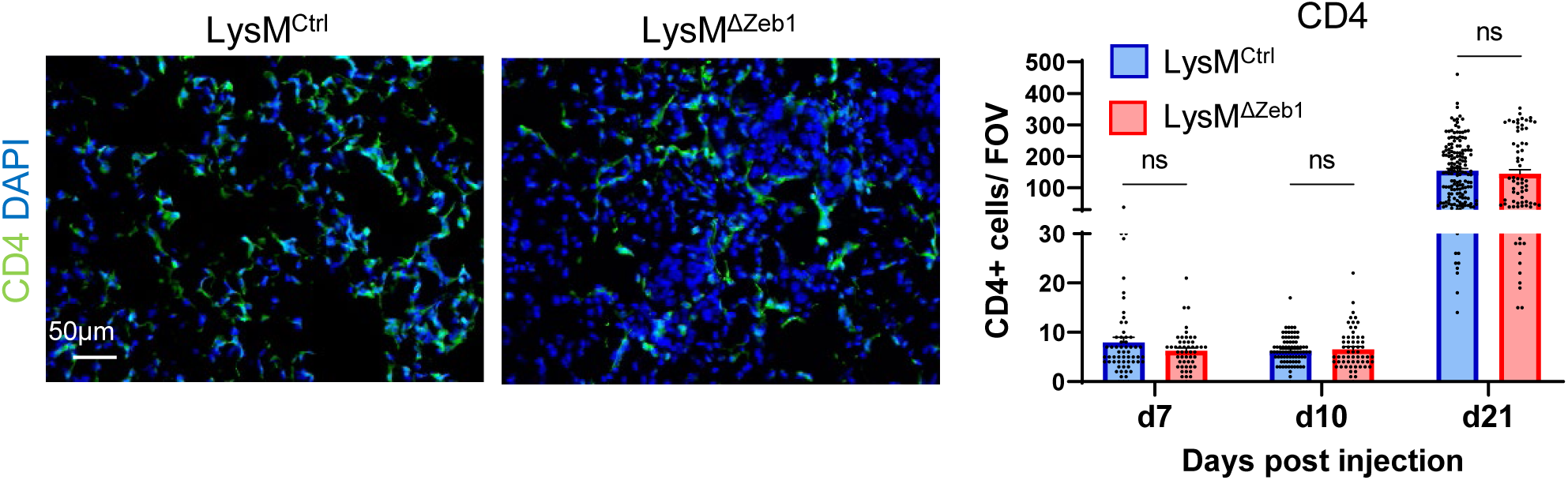
Infiltration of CD4+ cells into lung colonies is unaffected by loss of ZEB1 in macrophages. Representative images at 10 dpi and quantification over time of IF for CD4+ cells of KPC lung colonies in LysM^Ctrl^ and LysM^ΔZeb1^ mice with DAPI-stained nuclei (n>50 images; means +SEM; 2-way ANOVA).

