## Supplemental Table 3 for "Macrophages foster adaptive anti-tumor immunity by ZEB1-dependent cytotoxic T cell chemoattraction"

### Fuchs et al. Table S3 Material

#### Oligoucleotides

Primers for polymerase chain reaction (PCR)

| | Forward primer | Reverse primer |
|---|---|---|
| LysM-Cre | cttgggctgccagaatttctc | agcgattagctggagccatcaag<br>cccagaaatgccagattacg |
| Zeb1 del | cgtgatggagccagaatctgacccc | gccatctcaccagcccttactgtgc |
| Zeb1 flox | cgtgatggagccagaatctgacccc | gccctgtctttctcagcagtgtgg<br>gccatctcaccagcccttactgtgc |

Primers and UPL numbers for quantitative reverse transcriptase PCR (qRT-PCR)

| | Forward primer | Reverse primer | UPL # |
|---|---|---|---|
| Gapdh | agcttgtcatcaacgggaag | tttgatgttagtggggtctcg | 9 |
| Snai1 | cttgtgtctgcacgacctgt | caggagaatggcttctcacc | 71 |
| Snai2 | cattgccttgtgtctgcaag | agaaaggcttttccccagtg | 71 |
| Twist | agctacgccttctccgtct | tccttctctggaaacaatgaca | 58 |
| Zeb1 | aggtgatccagccaaacg | ggtggcgtggagtcagag | 93 |
| Zeb2 | ccagaggaaacaaggatttcag | aggcctgacatgtagtcttgtg | 42 |

#### Antibodies

Primary antibodies

| Specificity | Host | Catalog no. | Manufacturer | Dilution |
|---|---|---|---|---|
| α-Tubulin | mouse | T6199 | Sigma-Aldrich | 1:5000 |
| CD8 | rabbit | 50389-T26 | Sino Biologicals | 1:500 |
| CD68 (human) | mouse | ab201973 | Abcam | 1:100 |
| CD68 | rabbit | PA5-78996 | Invitrogen | 1:600 |
| cleaved Caspase-3 | rabbit | 9664S | Cell Signaling | 1:200 |
| F4/80 | rat | MCA4976 | Bio-Rad | 1:200 |
| ZEB1 | rabbit | HPA027524 | Sigma-Aldrich | 1:1000 (IHC)<br>1:2000 (WB) |
| ZEB1 | rabbit | NBP1-05987 | Novus Biologicals | 1:250 |
| ZEB1 | rabbit | E2G6Y #70512 | Cell Signaling | 1:400 (IF) |
| ZEB1 | mouse | AMAb90510 | Sigma-Aldrich | 1:300 |

Secondary antibodies

| Specificity | Host | Catalog no. | Manufacturer | Dilution |
|---|---|---|---|---|
| AlexaFluor488 anti-rabbit IgG (H+L) | goat | A11034 | Sigma-Aldrich | 1:200 |
| anti-rabbit-HRP polymer | goat | K4003 | DAKO | 1:1 |
| anti-rat-HRP | rabbit | A18915 | life technologies | 1:500 |
| CF640R anti-rabbit IgG (H+L) | goat | SAB4600164 | Sigma-Aldrich | 1:200 |
| CF640R anti-mouse IgG (H+L) | goat | SAB4600343 | Sigma-Aldrich | 1:200 |
| anti-rabbit-HRP (WB) | goat | 111-035-144 | Dianova | 1:10000 |
| anti-mouse-HRP (WB) | goat | 115-035-146 | Dianova | 1:10000 |

FACS antibodies

| Specificity | Catalog no. | Manufacturer | Dilution |
|---|---|---|---|
| TruStain FcX™ PLUS (anti-mouse CD16/32) antibody | 156603 | BioLegend | 1:200 |
| Purified anti-mouse CD16.2 Clone 9E9 | 149502 | BioLegend | 1:400 |
| InVivoMAb anti-mouse CD16/CD32 Clone 2.4G2 | BE0307 | Bio X Cell | 1:400 |
| B220-APC | 103211 | BioLegend | 1:200 |
| CCR3-APC | 144511 | BioLegend | 1:200 |
| CD3e-Biotin | 100304 | BioLegend | 1:200 |
| CD3e-Al647 | 100322 | BioLegend | 1:200 |
| CD4-Al488 | 100532 | BioLegend | 1:50 |
| CD4-APC-Fire750 | 100568 | BioLegend | 1:400 |
| CD4-BV605 | 100547 | BioLegend | 1:200 |
| CD8a-Al647 | 100727 | BioLegend | 1:200 |
| CD8a-BV570 | 100740 | BioLegend | 1:400 |
| CD11b-Al700 | 101222 | BioLegend | 1:400 |
| CD11b-BV421 | 101235 | BioLegend | 1:150 |
| CD11c-BV510 | 117337 | BioLegend | 1:100 |
| CD11c-PE | 117307 | BioLegend | 1:100 |
| CD11c-PE-CF594 | 562454 | BD Biosciences | 1:400 |
| CD19-Biotin | 115504 | BioLegend | 1:400 |
| CD19-BUV737 | 612781 | BD Biosciences | 1:400 |
| CD25-PE | 102007 | BioLegend | 1:200 |
| CD45-APC | 103111 | BioLegend | 1:200 |
| CD45-BV605 | 563053 | BD Biosciences | 1:400 |
| CD45-FITC | 103107 | BioLegend | 1:200 |
| CD45R (B220)-PE-Cy5 | 103210 | BioLegend | 1:800 |
| CD49b (DX5)-PE-Cy5 | 15-5971-82 | eBioscience | 1:200 |

| | | | |
|---|---|---|---|
| CD49R (B220)-PE-CF594 | 562290 | BD Biosciences | 1:800 |
| CD68-FITC | 137006 | BioLegend | 1:200 |
| CD88-PerCP-Cy5.5 | 135813 | BioLegend | 1:200 |
| CD90.2-Biotin | 105304 | BioLegend | 1:400 |
| CD103-BV711 | 121435 | BioLegend | 1:400 |
| CD115-PE | 135505 | BioLegend | 1:100 |
| CD161b/ c-BUV395 | 564144 | BD Biosciences | 1:400 |
| CD326-BUV737 | 741818 | BD Biosciences | 1:400 |
| CX3CR1-BV711 | 149031 | BioLegend | 1:400 |
| EPCAM-BV510 | 118231 | BioLegend | 1:200 |
| F4/80-Al647 | 123122 | BioLegend | 1:400 |
| F4/80-PERCP-Cy5.5 | 123127 | BioLegend | 1:100 |
| LY-6C-APC | 128015 | BioLegend | 1:200 |
| LY-6C-APC-e780 | 47-5932-82 | eBioscience | 1:200 |
| LY-6G-BV510 | 127633 | BioLegend | 1:100 |
| LY-6G-V450 | 562366 | BD Biosciences | 1:200 |
| MerTK-PE-Cy7 | 25-5751-82 | eBioscience | 1:100 |
| MHC II-PE-Cy7 | 107629 | BioLegend | 1:200 |
| MHC II-V500 | 562366 | BD Biosciences | 1:200 |
| NK1.1-PE-Cy7 | 108713 | BioLegend | 1:200 |
| Streptavidin-BUV496 | 612961 | BD Biosciences | 1:50 |
| Siglec F-FITC | 155503 | BioLegend | 1:100 |
| TCRβ-PE | 109207 | BioLegend | 1:100 |
| Ter119-Biotin | 116204 | BioLegend | 1:800 |
| XCR1-BV650 | 148220 | BioLegend | 1:400 |
| Zombie-NIR™ Dye | 77184 | BioLegend | 1:750 |
